# Two-photon activation, deactivation, and coherent control of melanopsin in live cells

**DOI:** 10.1101/2025.03.26.645437

**Authors:** Carlos A. Renteria, Jiho Kahng, Brian Tibble, Rishyashring R. Iyer, Jindou Shi, Haya Algrain, Eric J. Chaney, Edita Aksamitiene, Yuan-Zhi Liu, Phyllis Robinson, Tiffany Schmidt, Stephen A. Boppart

## Abstract

Intrinsically photosensitive retinal ganglion cells are photoreceptors discovered in the last 20 years. These cells project to the suprachiasmatic nucleus of the brain to drive circadian rhythms, regulated by ambient light levels. The photopigment responsible for photoactivation in these cells, melanopsin, has been shown to exhibit many unique activation features among opsins. Notably, the photopigment can exist in three states dependent on the intensity and spectrum of ambient light, which affects its function. Despite increasing knowledge about these cells and melanopsin, tools that can manipulate their three states, and do so with single-cell precision, are limited. This reduces the extent to which circuit-level phenomena, and studying the implications of melanopsin tri-stability in living systems, can be pursued. In this report, we evoke and modulate calcium transients in live cells and intrinsically photosensitive retinal ganglion cells from isolated retinal tissues following two-photon excitation using near-infrared light pulses. We demonstrate that two-photon activation of melanopsin can successfully stimulate melanopsin-expressing cells with high spatio-temporal precision. Moreover, we demonstrate that the functional tri-stability of the photopigment can be interrogated by multiphoton excitation using spectral-temporal modulation of a broadband, ultrafast laser source.

## Introduction

Intrinsically photosensitive retinal ganglion cells (ipRGCs) are increasingly studied since their initial discovery over two decades ago.^1,2^ These cells and their many subtypes^3,4^ have been implicated in circadian rhythm regulation^5–10^, largely due to their projection to the suprachiasmatic nucleus (SCN) of the brain, the region responsible for this regulation.^11–14^ The photopigment, melanopsin (OPN4), has also been discovered to have a number of unique properties that support regular ipRGC function. These include much slower activation and decay kinetics relative to rods and cones^15,16^, a prolonged and persistent photoresponse^3,15,17,18^, and most interestingly, the ability to switch between three different functional states based on the wavelength they are illuminated with^19–21^. These features make for a unique, naturally-occurring photopigment that robustly integrates ambient light to drive circadian rhythms, regulate the pupillary light reflex^22^, and many other non-image forming functions.

Melanopsin, the photopigment in ipRGCs, has been shown to have slower activation kinetics^15,19^ (∼1s) compared to rhodopsin and cone opsins (∼108 ms)^23^. Melanopsin-expressing cells also have substantially longer activation periods, on the order of several seconds, compared to rhodopsin-expressing cells, which last a few milliseconds. Whereas rods and cones in the retina photobleach immediately upon activation and take up to 30 minutes to recover, melanopsin-expressing cells have higher tolerance to photobleaching^24^. Most interestingly, melanopsin is a tri-stable protein with possible photoactivated conversions between three retinal isomers-all-trans retinal (metamelanopsin) |*M*⟩, 11-cis retinal (melanopsin) |*R*⟩, and 7-cis retinal (extramelanopsin) |*E*⟩, illustrated in Supplementary Fig. 1 (detailed in Supplementary Note 1). The states are so named by historical convention, based on the isomerization state of the retinal photopigment^19,25^. In the single photon regime, excitation with blue light can induce a transition from |*R*⟩ to |*M*⟩ and from |*E*⟩ to |*M*⟩. When the photopigment is in the |*M*⟩ state, excitation with yellow light induces a transition from |*M*⟩ to |*E*⟩ state. Since both |*M*⟩ and |*E*⟩ are excited states of the photopigment, both have natural relaxation transitions to the ground state |*R*⟩, with different relaxation kinetics. The implications of this tri-stable nature is still an open question, though it has been proposed to facilitate uniform spectral activation, finer control over pigment function by control of its states, a broadened excitation spectrum, and long-term stabilization towards a photoequilibrium, that are all otherwise not features in bistable photopigments^19^.

While many of the absorptive properties of melanopsin are currently known, investigations into the functional circuitry of melanopsin-expressing cells, notably ipRGCs, are lacking. Studies that pursue ipRGC circuits do so primarily through morphological identification of projections from ipRGCs to the SCN^4,26^, and use genetic tools to tag these cells and their projections^4,15,27,28^ or ablate distinct ipRGCs to unveil possible downstream effects^4,7,8,29^. These tools thoroughly identify projections to the SCN, however, they are limited in that cellular function cannot be evoked or measured *in situ* at a large scale. Moreover, illumination paradigms rely on visible light illumination, which would be absorbed by all exposed ipRGCs due to their high sensitivity^15^. Thus, the effects of distinct subpopulations of ipRGCs on SCN signaling and other downstream effects cannot be resolved in their native environment. Moreover, how these features combine with the inherent tri-stable nature of melanopsin cannot be performed with existing tools, leaving the implications for melanopsin tri-stability an open question. Advances in light delivery with single-cell precision and state-to-state control are needed to elucidate these effects.

Two-photon absorption is a tool which has increasingly become adopted in neuroscience for stimulating highly-specific networks of cells with single-cell precision^30–34^, and has long been used for two-photon excited fluorescence microscopy^35–37^. This optical method can help address this challenge. Due to the low probability of a two-photon absorption event occurring within light-absorbing chromophores exposed to near-infrared (NIR) light^38,39^, absorption events are confined to a small focal region with subcellular resolution^38^. This permits generation of images with reduced out-of-focus excitation^38^, stimulation of photosensitive cells with single-cell precision^30– 33,40^, and increased depth penetration due to use of longer wavelengths^41^. Despite these advantages, two-photon absorption alone still comes with its own inherent limitations. Optimal two-photon absorption requires temporally short pulses to achieve, with optical systems and tissues adding dispersion that broadens pulse duration^34,36,38,42–44^. Many new lasers have internal dispersion compensation mechanisms, and dispersion is frequently compensated for using prism pairs^45^. However, for applications that require tuning laser wavelength, dispersion needs to be compensated for each wavelength, increasing system complexity, and increasing time between stimulations. This makes single-cell stimulation across multiple wavelengths with equal efficiency challenging, requiring engineering efforts to realize this in melanopsin-expressing cells.

In this work, we demonstrate successful two-photon stimulation of individual melanopsin-expressing HEK293T cells (ATCC, CRL 3216), and ipRGCs *in situ* in isolated murine retina using calcium imaging to monitor cellular activation. Moreover, we demonstrate that chromatic control of melanopsin in HEK293T cells can be leveraged by spectral-temporal modulation of our supercontinuum source^34,46^ to promote activation and deactivation of melanopsin-expressing cells in the two-photon regime. We do so using a two-photon microscopy system (Supplementary Figs. 2-5) that leverages a supercontinuum of light (900-1160 nm) with ultrashort, high peak power pulses (20 MHz repetition rate, ∼20 fs pulse width) to capture the broad absorption range of melanopsin^2,19^ for two-photon stimulation of its various states. The system was designed to simultaneously capture calcium imaging data, using the calcium indicator Calbryte™-590 AM (AAT Bioquest)^47^. This study establishes a precedent for two-photon stimulation experiments in melanopsin-expressing cells, and enables further investigations of ipRGC circuitry with single-cell precision.

## Results

### Two-photon activation of melanopsin-expressing HEK293T cells

To determine the effects of two-photon absorption on activation of melanopsin-expressing cells, we transfected human embryonic kidney cells (HEK293T, ATCC) with a plasmid encoding melanopsin with a GFP tag^17^. The cells were also loaded with 5 µM Calbryte-590 AM to monitor transient changes in intracellular calcium following irradiation with a pulsed NIR stimulus. The experimental paradigm implemented here, using one laser for imaging Calbryte-590 AM, and another for stimulating melanopsin with different scan durations, is illustrated in Fig. 1A. Successful transfection permits identification of melanopsin-expressing cells with the presence of GFP (Fig 1B, green, and Supplementary Fig. 6). The calcium indicator was indiscriminately loaded and identifiable in all cells (Fig 1B, magenta). The model was validated by irradiation with 488 nm light and widefield calcium imaging (Supplementary Fig. 7) to test the single-photon responses, which exhibited kinetics consistent with previous studies^17^. Following irradiation with pulsed NIR light (1 s scan duration using a laser with 20 MHz repetition rate, ∼20 fs pulse duration, and wavelengths spanning 900-1160 nm, see Methods) scanned across a single melanopsin-expressing HEK293T cell, a rapid and prolonged increase in calcium levels was elicited only in the targeted (Fig. 1C, Cell 1). Following subsequent stimulations, consistent calcium responses were evoked from that individual cell that were not present in adjacent, non-targeted GFP^+^ cells (Fig. 1C, Cells 2 and 3) or GFP^-^ cells (Fig. 1C, Cell 4). At these conditions, with 20 mW incident optical power and a 1 s scan duration, the Ca^2+^ responses had rise times of ∼4 s and decay times of ∼25 s (Fig. 1E), typical for melanopsin cell responses^17,18^ and indicative of successful single-cell, two-photon activation.

**Fig. 1:**
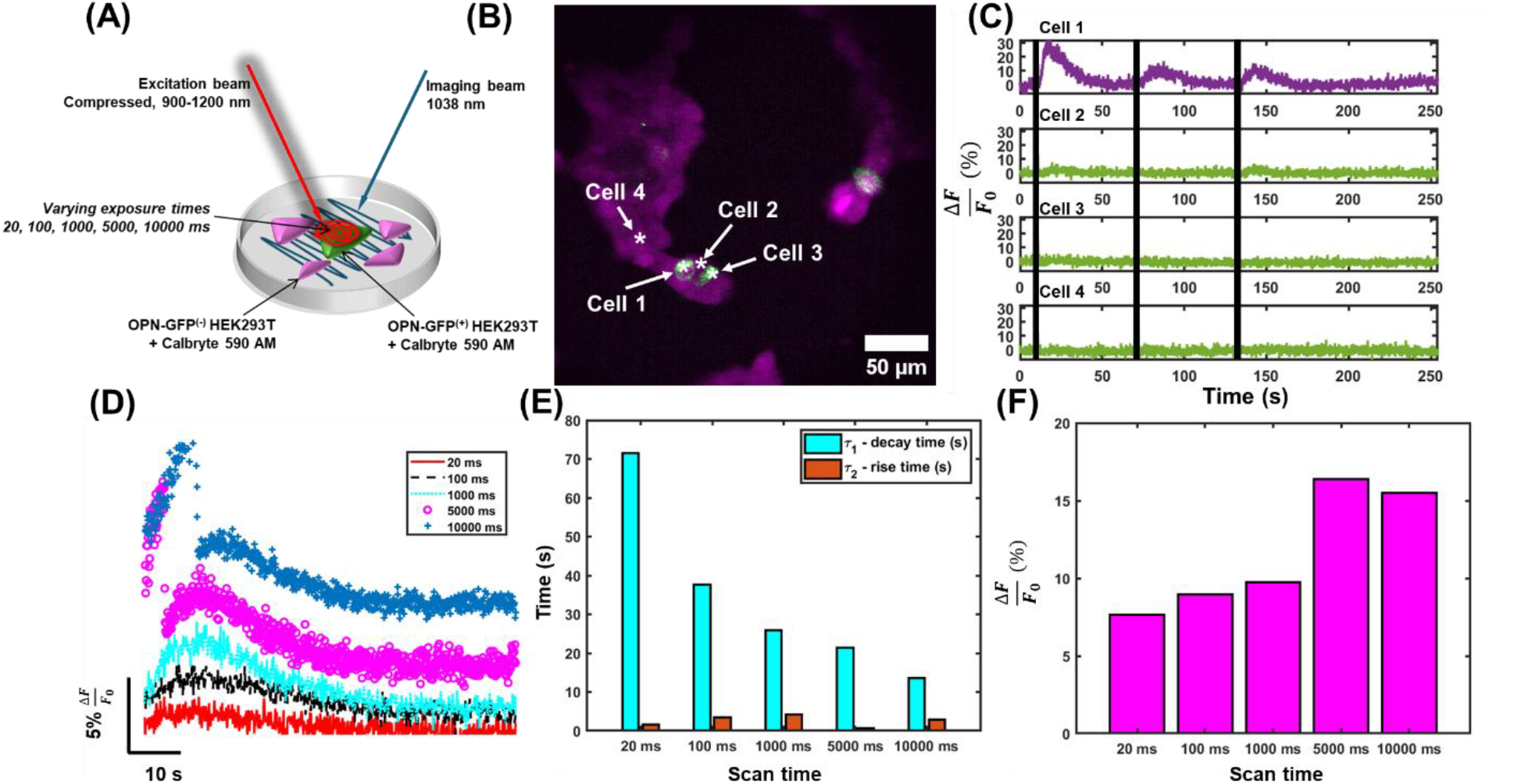
Activation kinetics following different scan durations on HEK293T cells. **a**, Illustration of the dual-laser scanning paradigm. The workflow consisted of a raster-scanned beam for calcium imaging and a spiral-scanned beam with variable scan duration for melanopsin-activation. **b**, Representative field-of-view of cultured HEK293T cells loaded with Calbryte-590 (magenta) and tagged with OPN4-GFP (green). **c**, Representative calcium transient from cells identified in A. Time of stimulation is denoted by black tick marks. **d**, Mean of all evoked calcium transients following irradiation with a supercontinuum for different amounts of time. **e**, The rise and decay constants from exponential fits applied to the data in **d. f**, The peak calcium response obtained from the traces in **d**. n = 52 calcium transients from N = 4 stimulated cells.

Next, the effects of the scan duration and stimulation energy on the Ca^2+^ amplitude and rise and fall times were explored. By varying the scan duration (as illustrated in Fig. 1A), we can evoke calcium transients with amplitudes that track with irradiation duration (Fig. 1D, F, Supplementary Fig. 8A). After fitting an average of several calcium traces for each illumination paradigm to a biexponential decay curve (see Methods), shorter decay times result, following increased irradiation levels (Fig. 1E, Supplementary Fig. 8B). This was found along with slightly increasing rise times between 20 and 1000 ms scan durations (Fig. 1E, Supplementary Fig. 8C), although the trends in rise times are less pronounced. Increased amplitudes of transient calcium responses were also associated with increased irradiation times (Fig. 1F). These trends are in contrast to that of other opsins, such as Channelrhodopsin-2 (ChR2) which have larger evoked responses when irradiated with faster scan durations^40^. This is largely due to the integrative nature of melanopsin.

Melanopsin absorbs photons across different wavelengths, space, and time to promote continued excitation in ipRGCs^15^, consistent with its role in regulating excitation based on ambient lighting. Biophysically, melanopsin also has deactivation kinetics that are two orders of magnitude slower than that of ChR2^15,19,48^, and irradiation times to cover the full surface of a single opsin-expressing cell need only be faster or equal to the decay time of the opsin^40^. This makes melanopsin-expressing cells uniquely capable of absorbing many more photons at one time compared to their other opsin-expressing cells.

This integrative behavior is also apparent when melanopsin is irradiated with NIR light at increasing powers (as illustrated in Fig. 2A, with representative sample Fig. 2B), showing increased amplitude of calcium responses when illumination power is varied between 5 mW and 30 mW (Fig. 2C,F, Supplementary Fig. 9) with the scan duration fixed at 1 s. Fitting the mean traces for each illumination power (Fig. 2D) shows decreased decay times and slightly increased rise times with increasing illumination power (Fig. 2E, Supplementary Fig. 9B,C) accompanied by increased magnitudes of calcium responses for average traces (Fig. 2E) and all individual stimulation epochs (Supplementary Fig. 9A). These are consistent with the results for increased scan durations in Fig. 1. Together, these results show that increased irradiation over time leads to increases in the amplitude of calcium signals, decreases in decay times, and slight increases in rise times from increased numbers of absorbed photons (Supplementary Fig. 10), similar to melanopsin’s spatiotemporal photon integration properties in the single-photon regime^48^. We show that the absorptive nature of melanopsin in the two-photon regime is consistent with reported dynamics in the single-photon regime^15^.

**Fig. 2:**
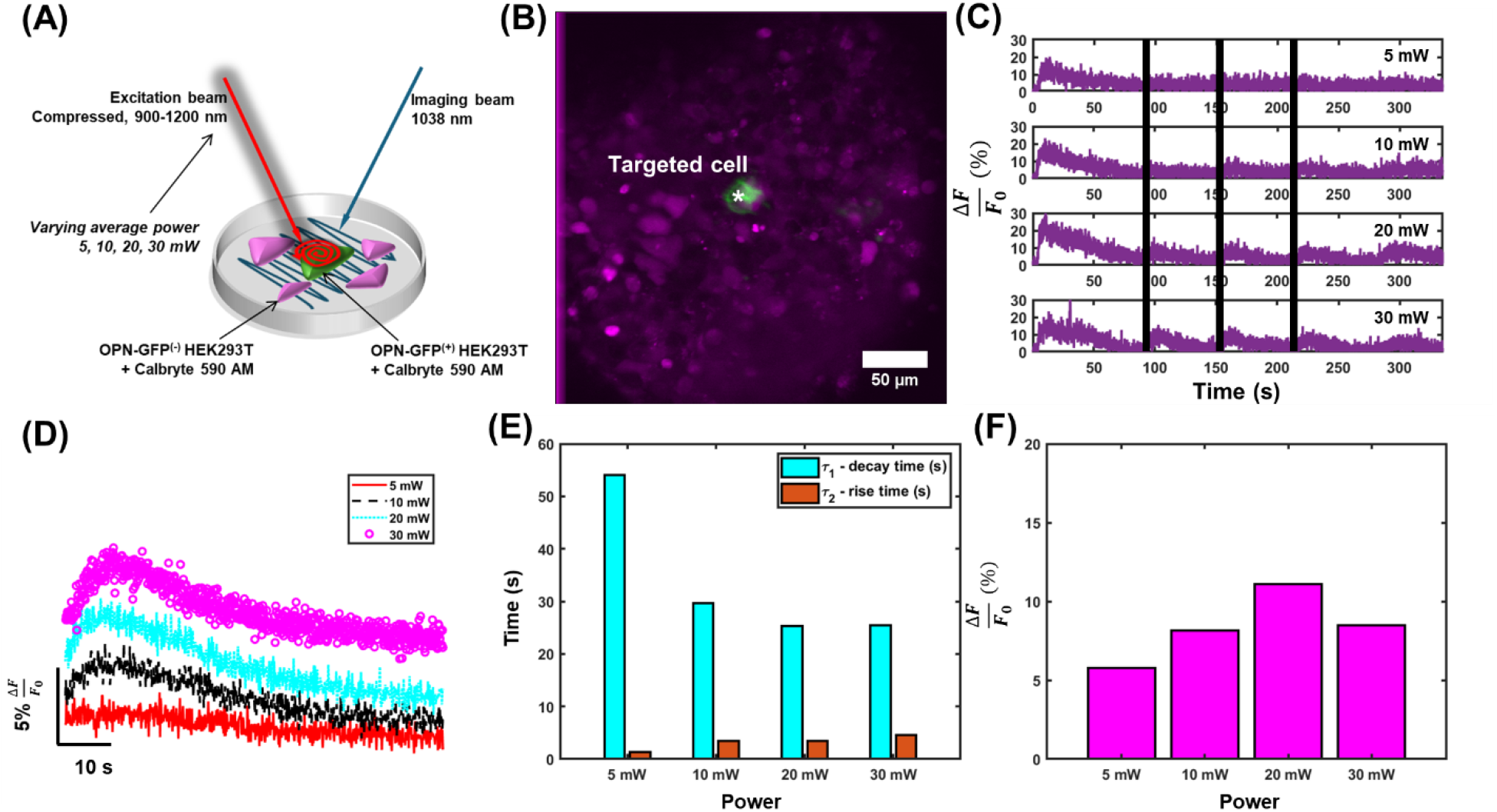
Activation kinetics following different average powers irradiated on HEK293T cells. **a**, Illustration of the dual-laser scanning paradigm. The design consisted of a raster-scanned beam for calcium imaging and a spiral-scanned beam with variable incident power for melanopsin-activation. **b**, Representative field-of-view of cultured HEK293T cells loaded with Calbryte-590 (magenta) and tagged with OPN4-GFP (green). **c**, Representative calcium transient from cells identified in **b**. Time of stimulation is denoted by vertical black bars. **d**, Mean of all evoked calcium transients following irradiation with a supercontinuum for different amounts of time. **e**, The rise and decay constants from exponential fits applied to the data in **d. f**, The peak calcium response obtained from the traces in **c**. n = 188 calcium transients from N = 9 cells.

### Chromatic distinction and coherent control of melanopsin in OPN4-GFP HEK293T cells

Melanopsin can exist in three different states based on irradiation wavelength^25^, which causes activation or deactivation of the opsin, and consequently, changes the cellular activity^19,21,49^. We leveraged the supercontinuum generated in this work (Supplementary Figs. 3-5) to probe the activation kinetics in three different illumination ranges in OPN4-GFP HEK293T cells to determine if this phenomena can be evoked with two-photon absorption. Cultured cells were illuminated with NIR light between 900-1000 nm to permit activation of the cell (*via* transition of OPN4 from |*R*⟩ to |*M*⟩ and |*E*⟩ to |*M*⟩), 1100-1200 nm to permit deactivation of the cell (*via* transition of OPN4 from |*M*⟩ to |*E*⟩), and the full supercontinuum of light (900-1200 nm), which should also primarily evoke excitation to the |*M*⟩ state based on the response of melanopsin to white light sources in the single-photon regime^19^.

We leveraged a rapid raster scanning approach, where the supercontinuum was used to simultaneously drive melanopsin excitation and capture image data (Fig. 3A) at 1.63 Hz, faster than the deactivation kinetics of melanopsin^15^. When melanopsin is activated during illumination, a sustained Ca^2+^ response is expected^17,18^. During deactivation, the Ca^2+^ responses will not be prolonged or sustained^21^. Representative fields-of-view (Fig. 3B) and traces (Fig. 3C) show co-expression of these fluorophores in culture (Fig. 3B). The representative calcium transients evoked from OPN4^+^ are sustained during illumination (Fig. 3C, Supplementary Fig. 11, Cells 1-3) and are absent in OPN4^-^ cells (Supplementary Fig. 11, Cells 4-6). For a representative field-of-view (Fig. 3D), different scan paradigms were implemented that involved a knife-edge filter for spectral control of melanopsin irradiation (Fig. 3E), and modification of the spectral-phase of the supercontinuum source (Fig. 3F) for predictably driving melanopsin to a desired final state during imaging (see Methods and Supplementary Note 2 for full details). Following illumination of a region of interest containing an OPN4^+^ cell (Fig. 3D) with the full supercontinuum (Fig. 3G, left) and spectra between 900-1000 nm (Fig. 3G, middle), a sustained calcium response throughout image capture and irradiation was evoked. Following illumination with wavelengths between 1100-1200 nm (Fig. 3G, right), the same cell shows a complete drop in calcium levels to well below that of the original sustained response, indicative of a net transition to its inactivate state (the full photocycle of melanopsin is described in the Supplementary Note 1).

**Fig. 3:**
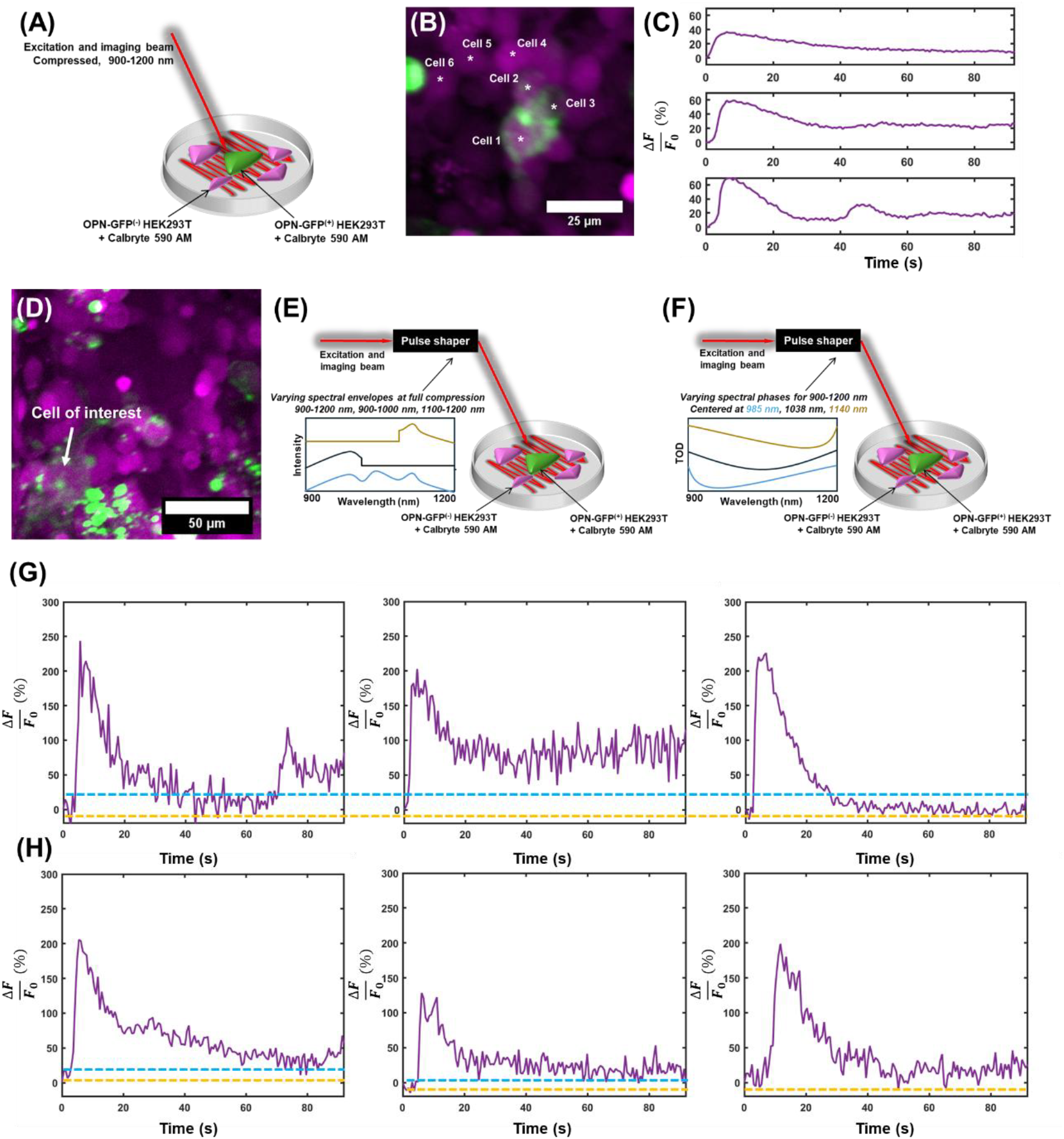
Evaluating the effects of the spectral properties of the supercontinuum on evoked calcium responses. **a**, An illustration of the scanning paradigm. A single laser consisting of a supercontinuum of light was raster-scanned across a field of cells, simultaneously evoking fluorescence from the calcium indicator and activation melanopsin in transfected cells. **b**, Representative field-of-view for these experiments. **c**, Representative calcium transients from activated, melanopsin-expressing cells in **b. d**, Representative field-of-view for results in **g, h. e**, Illustration of the single-beam paradigm used for results in **g**, with the spectrum controlled by blocking portions of the spectrum. **f**, Illustration of the single-beam scanning paradigm for results in **h**, where TOD is used to drive melanopsin activation by coherent control (see Supplementary Note 2). **g**, Calcium responses from irradiated HEK293T cells illuminated with the full supercontinuum (left), 900-1000 nm (middle), and 1100-1200 nm (right) from the supercontinuum of light. **h**, Effects of TOD tuning on evoked calcium responses. Notably, after tuning the central wavelength to <980 nm (left), ∼1030 nm (middle), and >1140 nm (right).

In the multiphoton regime, researchers have demonstrated that the spectral phase of ultrafast laser pulses can be used to predictably drive photochemical reactions^43,44,50–55^, a principle known as coherent control (further detailed in the Supplementary Note 2). This principle has also been used to modify the yield of opsins^56,57^, and modify activation kinetics in opsin-expressing neurons^42^. To this end, we modulated the amount and the central frequency of third-order dispersion (TOD) of the supercontinuum to predictably control fluorescence absorption^55,58^ to drive melanopsin to activation or deactivation (further details in the Supplementary Note 2). When the central wavelength of the TOD is changed from 980 nm (activation phase, Fig. 3H, left) to 1030 nm (central wavelength phase, Fig. 3H, middle) to 1140 nm (deactivation phase, Fig. 3H, right), the Ca^2+^ responses of the cells change as expected. A sustained activation is observed when the TOD is tuned to the central wavelength and below, with a gradual decrease in the sustained response as the central wavelengths of the TOD are shifted to longer wavelengths. Thus, the activity of OPN^+^ cells could be modulated using two-photon absorption with the same source using identical spectral envelopes but with varying spectral phase profiles.

To determine if melanopsin can be dynamically activated and deactivated, additional experiments were performed where the spectral phase was tuned to either activation or deactivation of melanopsin initially, and the phase later reversed during imaging (Fig. 4A; spectral phase diagram magnified in Fig. 4B). For representative cells (Fig. 4C), when activation pulses preceded the deactivation pulses, few additional calcium responses were evoked (Fig. 4D, G) consistent with the anticipated prolonged reversal of melanopsin inactivation to the |*E*⟩ state. When deactivation pulses preceded activation pulses, more robust calcium responses were evoked in the representative cells (Fig. 4E, H), indicative of a transition of melanopsin activation to the |*M*⟩ state, and consequent increases in calcium levels. In cells that did not show a strong calcium transient, a larger calcium baseline during the sustained response period was occasionally apparent (Fig. 4G and Supplementary Fig. 11B,C), which could still indicate a slight increase in the population of melanopsin in its active |*M*⟩ state, but not sufficient enough for a characteristic calcium response in the cell. Because of the broad absorption spectrum of melanopsin in each state^19^, it is possible for many of the melanopsin molecules to exist in some equilibrium between states, which can affect the net evoked calcium transients in individual cells when these transitions are invoked. Nonetheless, these results demonstrate that the coherent control principle can be leveraged in melanopsin-expressing cells to drive activation kinetics towards one of the two desired light-evoked states, increasing the dimensionality of optical tools available for two-photon activation of melanopsin-expressing cells. This is corroborated with other cells, shown in the representative traces in Supplementary Fig. 11.

**Fig. 4:**
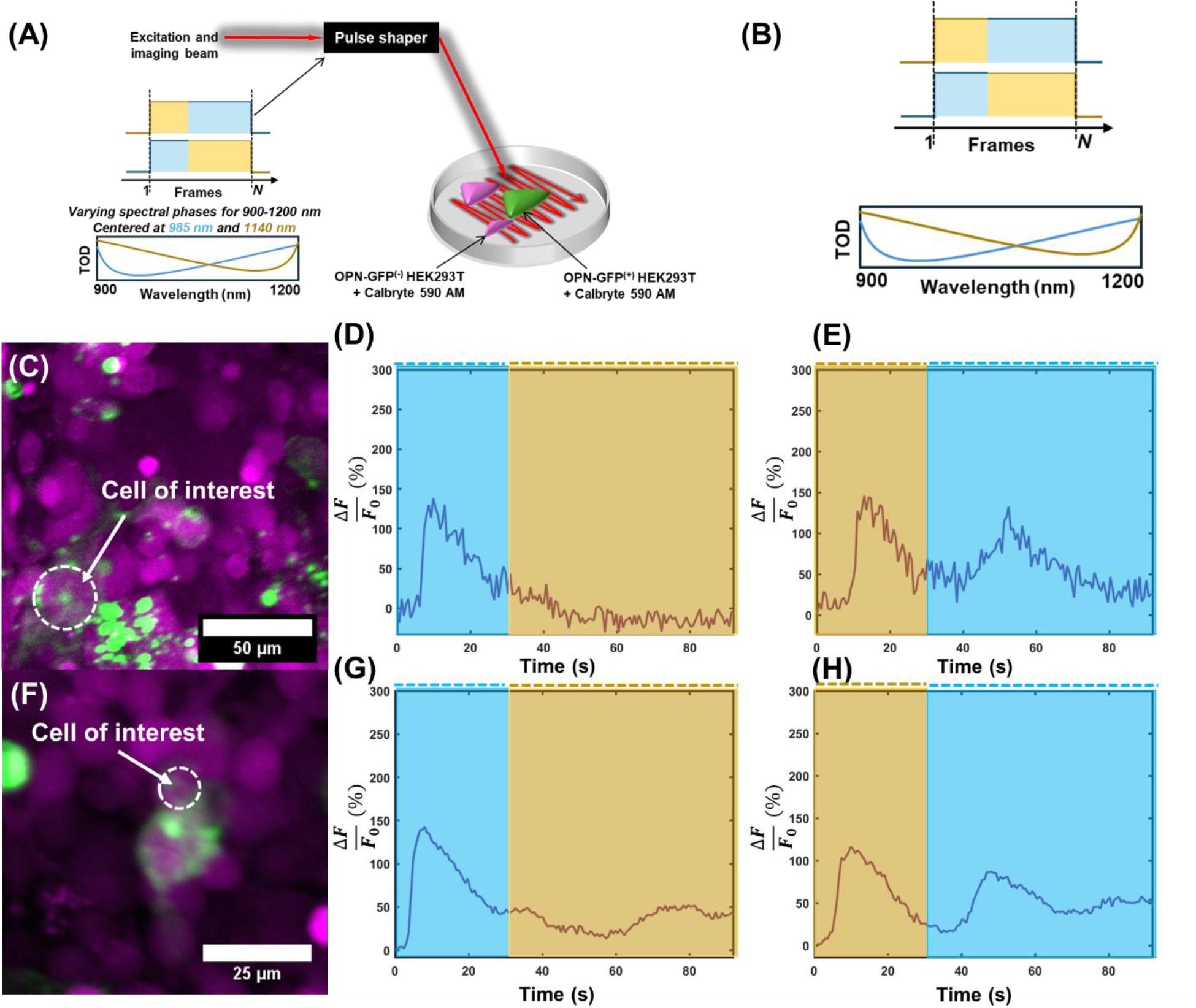
Dynamic evaluation of coherent control of melanopsin in HEK293T cells. **a**, Illustration of the dynamic experimental paradigm. A single supercontinuum beam was raster-scanned across different cells. The TOD was tuned towards <980 nm 30 seconds, and immediately tuned to >1100 nm, or vice versa. Reversal from <980 nm to >1100 nm guides melanopsin from activation to deactivation at ∼30 s, whereas from >1100 nm to <980 nm causes reactivation of melanopsins in an inactive state (see Supplementary Notes 1 and 2) at ∼30 s. **b**, Inset from **a** showing the change in illumination during the experiment (top) and the corresponding phase profiles (bottom) for activation (blue) and inactivation (yellow) of melanopsin. **c**, Representative field-of-view. **d**, Evoked calcium response following initial irradiation with light tuned to 980 nm (blue), and tuned to 1140 nm (yellow) 30 seconds after imaging. **e**, Same as **d**, but with the reversed order. **f**, Another representative field-of-view. **g**, Evoked calcium response following initial irradiation with light tuned to 980 nm (blue), and tuned to 1100 nm (yellow) 30 seconds after imaging for the targeted cell in F. **h**, Same as **g**, with the reversed order.

### Two-photon in-situ activation of ipRGCs

Two-photon evoked calcium responses of ipRGCs were generated, following irradiation with NIR light. The same raster scanning paradigm used for simultaneous stimulation of melanopsin and imaging of Calbryte-590 AM in the previous section was used here. Isolated retina from adult mice whose ipRGCs express GFP^27^ were bulk-loaded with Calbryte-590 AM for calcium imaging (see Methods). The isolated retina was then placed under the microscope for identification of ipRGCs in the green GFP channel. Once identified (Fig. 5A, E) and baseline images captured, time-series image data was captured at ∼1.63 Hz to track changes in calcium levels using Calbryte-590 AM in the red channels. Evoked transients showed a brief, immediate increase in calcium level during the first few frames (Supplementary Fig. 12), followed by a rapid exponential decay in calcium levels throughout illumination. One instance is illustrated in Fig. 5B, Cell 1, and those from additional ipRGCs in Fig. 5F, Cells 1-3, and Cell 5. The videos used to generate the Fig. 5 plots are shown in Supplementary Vids. 1 and 2. The decays are reminiscent of those reported in ipRGCs loaded with Calbryte-590 AM that were stimulated with visible light, but imaged with two-photon microscopy at very low powers^59^. That is, following illumination and absorption of visible photons, the results in Caval-Holme *et. al*.^59^ show an immediate increase in calcium responses, followed by an exponential decay in calcium levels during sustained illumination^59,60^. This decay, found across all activated ipRGCs recorded in this study, is consistently absent in adjacent, randomly selected eGFP^-^ RGCs and surrounding retinal tissue (Fig. 5B, Cells 2-3 and Fig. 5F, Cell 4), demonstrating specificity to ipRGCs. Similar to the reported calcium trace results in HEK293T cells, an exponential decay can be fitted to the calcium decays to quantify the decay time of melanopsin in ipRGCs (see Methods), along with the magnitude of these results. The distribution of these values is shown in Fig. 5D, with an average ΔF/F of ∼5.4%, and decay time of ∼3.2 s, within seconds of the decay times reported here for the HEK293T cells. Additional analysis of the decays also shows comparable peak ΔF values between conditions (Fig. 5G), but statistically significant decreases in the magnitude of the two-photon evoked calcium responses in active ipRGCs, notably the last ∼1 second of the recordings, compared to inactive cells (Fig. 5H), further showing specificity towards ipRGCs, and thus evoked two-photon responses from these cells. This is largely because inactive cells had flat calcium traces, compared to the exponential decays from ipRGC activation reported here. These data support two-photon absorption evoked responses in ipRGCs, extending the feasibility of this nonlinear optical phenomena in native melanopsin-expressing murine ipRGCs.

**Fig. 5:**
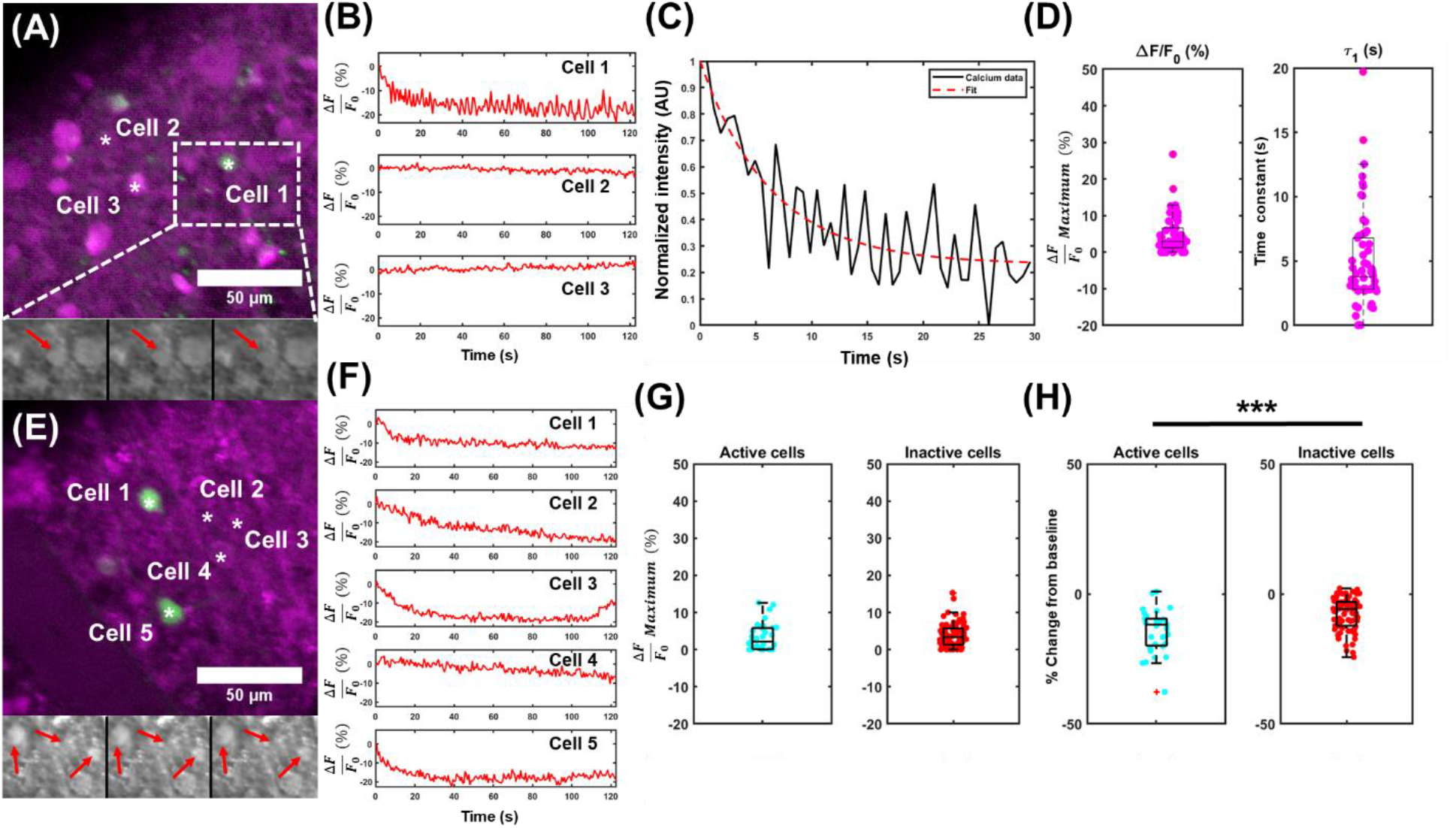
Two-photon absorption in ipRGCs. **a**, Representative field-of-view from isolated retina loaded with Calbryte-590 (magenta) with GFP-tagged ipRGCs (green). Multiple frames are illustrated below to show the changes in ipRGC calcium intensity over time. **b**, Calcium traces from a subset of cells identified in **a. c**, Representative monoexponential fit to the calcium decay for Cell 1 illustrated in **b. d**, Peak calcium transient (left) and the exponential decay constant (right) obtained by fitting a monoexponential decay to the time-series calcium traces for all ipRGCs identified in this study. **e**, Another representative field-of-view, with time-series changes shown below in gray. **f**, Additional calcium transients, and characteristic exponential decay shown for ipRGCs (Cells 1, 2, 3, and 5), along with an inactive ipRGC (Cell 4). **g**, Peak ΔF/F_0_ for all active ipRGCs and a subset of inactive cells identified in this study. **h**, The ΔF/F_0_ value calculated for the last ten frames for all identified active (left) and randomly chosen subset of inactive (right) cells in this study. n = 34 calcium transients identified from N = 34 cells in 6 retinae, p = 1.84*10^−5^.

## Discussion

The reported research advances provide a benchmark for further investigations that aim to directly probe functional ipRGC circuitry in the retina. Since ipRGCs have been shown to affect a diversity of circadian functions, including sleep, thermoregulation, and other functions, such tools can be used to more comprehensively evaluate what cells, and patterns of cells, affect this downstream regulation. Future work will be needed to determine the degree to which two-photon absorption can be leveraged in living organisms. To that end, developing an *in vivo* system for directed stimulation of ipRGCs and monitoring their calcium traces, as performed *in situ* in this work, will help realize direct observation of these functional circuits in living mice. Moreover, the ability to stimulate individual and combinations of cells can help resolve functional ipRGC circuits. By combining this capability with a spatial light modulator^31–33,61–63^ or other spatial patterning device for single-cell targeting, circuits implicated in various circadian rhythms can be further resolved. For example, functional circuits implicated in different circadian rhythms, such as sleep, thermoregulation, etc.^7,10^ can be disparately targeted and evaluated in living mice using two-photon excitation. Thus, evaluating how two-photon excitation affects different circadian behaviors can be explored through this enabling technique. The long decay kinetics of melanopsin further permit this, as illumination can be performed continuously for continued excitation, rather than having to rely on sequential stimulation due to rapid decay times typical in other opsins, such as ChR2^40^. This combination of factors makes melanopsin excitation an ideal candidate for continued investigation for single-cell, two-photon activation of functional ipRGC circuits.

This would be especially interesting to evaluate in the developing retina, which gives rise to spontaneous calcium waves^59,64,65^, and undergoes changes to melanopsin subtype population and melanopsin expression during development^4,66^. To-date, six different types of ipRGCs that differ in size, shape, dendritic branching, and melanopsin among other features, have been identified^4^. How each individual subtype functions in the ipRGC-SCN pathway, and how they regulate the different circadian rhythms, would be noteworthy to explore, though the M4-M6 subtypes, known for their lower melanopsin concentration and low GFP-expression^3,4,27,66^, may be difficult to identify. Combined advances in optics and genetic engineering of animal and cell lines for identifying and knocking-out ipRGCs^4,15,27,28,67^ may serve to facilitate this *in vivo*. Furthermore, how these circuits are differentially affected by different wavelengths of light, especially due to the tristable nature of melanopsin^19^, makes for a new unexplored field of study that can be realized using the technology and methodology introduced here.

On the topic of melanopsin tri-stability, few other techniques permit simultaneous activation of cells at the single-cell level with the ability treat them as a photoswitch with high efficiency. The unique nonlinear properties of two-photon absorption permit coherent control to be leveraged for this purpose^38,50,51,53,68,69^. Though a tunable laser source can be used for these same purposes if the central wavelength is tuned towards a spectral region favored by one melanopsin state, dispersion needs to simultaneously be accommodated for when laser wavelengths are switched. This requires a laser system with rapid wavelength tunability and known dispersion compensation, especially if optogenetic or *in vivo* applications ever require rapid photoswitching. Use of a pulse-shaper for coherent control especially provides a rapid mechanism for switching between states, as phase profiles can be applied instantaneously, whereas multi-wavelengths sources would require separate tunability and dispersion compensation to accomplish this task, though advances in rapid tunability could obviate this limitation. Coherent control, however, can be challenging, and requires a stable and well-tuned pulse-shaper to perform effectively^34,36,42,50–54,68,69^, so researchers should consider what is most appropriate for their systems when pursuing similar investigations.

Lastly, it has been reported that deactivation of melanopsin to the |*E*⟩ state requires more intense and long exposure to long wavelengths to adequately evoke in the visible-light regime^49^. Since two-photon absorption inherently uses high-intensity pulses for activation^34,36,38,42^, this technique can potentially resolve this limitation, and make applications involving melanopsin deactivation more easily realizable. As suggested with visible light, the combined capabilities of directed two-photon absorption of ipRGCs and chromatic control can be used as a new form of artificial light for mood disorders^49,70^. This could be used to develop new light-based therapies for ailments where circadian rhythm impairment has been implicated, including Alzheimer’s Disease^71^, jet-lag^72^, and seasonal affective disorder^72^, among others.

## Conclusion

We present here two-photon activation, deactivation, and coherent control of calcium transients in melanopsin-expressing HEK293T cells. This was demonstrated by illuminating melanopsin-expressing HEK293T cells with a supercontinuum of light for stimulation, illuminating cells with distinct spectral bands of the supercontinuum, and pulse-shaping of the supercontinuum to predictably drive melanopsin towards a desired final state. This was achieved using coherent control to drive deconstructive interference in undesired spectral bands of our supercontinuum. We also demonstrate that two-photon absorption can be used to evoke responses in ipRGCs from isolated murine retinae. The presented work establishes a framework for continued investigation into unexplored research paradigms in circadian and retinal science by bridging advances and novel engineering efforts in nonlinear microscopy^34,36–38,46,63^ and coherent control^50–53,55,58^ with biophysical and physiological knowledge of melanopsin kinetics^15,19,48,49^ in cultured cells^17,18,21^ and ipRGCs^3,4,59,60^.

## Methods

### System Design

The optical system was modified from a previously developed microscope^34^ and tailored to these specific experiments. The diagram is shown in Supplementary Figure 5. The laser source from a fiber-based laser (Satsuma, Amplitude Systemes) centered at 1030 nm was split into two using the combination of a Glan-Thompson polarizer and half-wave plate (HWP). One path was directed by a resonant scanner (RS) and galvanometer pair before being redirected to the microscope objective (XLPLN25XWMP2, Olympus) and then the sample plane. The second path was directed to a photonic-crystal fiber (PCF, LMA-PM-15, NKT Photonics) to generate a supercontinuum of light^36,73,74^. The supercontinuum was then directed to a pulse-shaper (MIIPS Box 640, Biophotonics Solutions) for phase-only modulation of the fiber supercontinuum. The amplitude of the supercontinuum was modulated by covering certain pixels of the SLM with a physical barrier. The supercontinuum was then directed to a galvanometer pair both for capturing images at the sample plane, or spiral scanning to illuminate a target cell. Fluorescence following illumination with either source is collected using photomultiplier tubes (PMTs). An analog PMT was used for high-speed imaging (H7422A-40, Hamamatsu) which was amplified through a transimpedance amplifier (TIA60, Thorlabs). A photon-counting PMT (H7421-40, Hamamatsu) was used for the alternative path. DIC imaging capabilities were integrated for identifying cells, using an EMCCD camera (QImaging, Retiga Electro) for detection, along with DIC filters.

### Pulse shaping

With the supercontinuum source, pulses were compressed to their near transform-limit using the multiphoton intrapulse interference phase scan (MIIPS) algorithm^68^, and verified using frequency-resolved optical gating (FROG)^75^. Additional phase profiles were imposed onto the SLM following compensation to their transform-limit. Experiments using third-order dispersion (TOD) applied a dispersion profile equivalent to *a*_3_(ω − ω_0_)^3^ to the SLM, where *a*_3_ is the third-order dispersion coefficient. In this work, we set *a*_3_ = 15,000 *fs*^3^, and ω_0_ to the anticipated peak for two-photon absorption of melanopsin. These wavelengths were ≤1000 nm for excitation, or ≥1100 nm in this work. The theory behind this choice was based on previous literature^50,55,58^ and simulations performed in this work which demonstrate that this combination of factors effectively maximizes two-photon absorption signals centered around ω_0_ while reducing contributions from undesired spectral windows (Supplementary Fig. 2). The effects of these and other phase profiles on ultrafast pulses and two-photon signals are shown in Supplementary Figs. 2 and 4.

### Cell culture preparation and transfection

HEK293T cells were grown on cell culture flasks (BioLite, 130192) with Dulbecco’s Modified Eagle’s Medium (DMEM) supplemented with 10% fetal bovine serum (FBS) and 1% Penicillin/Streptomycin in an incubator set to 37 ºC with 5% CO_2_ until ∼70-80% confluent. Thereafter, cell culture medium was removed, and cells were exposed to 0.25% trypsin for ∼1 minute. The trypsin was then removed, and cells were removed from the flask surface by gently pipetting DMEM over the trypsinized cells. Cultures were maintained by replating at a 1:5 ratio. Another 1 mL of cells was then transferred to a canonical tube for transfection. Transfection protocols were performed according to manufacturer-supplied procedures. Briefly, 100 µL of Opti-MEM I (Gibco, 31985062) was added to a separate container and incubated with ∼ 4 µg of melanopsin-GFP cloned in the mammalian expression plasmid^17,18^. Thereafter, 8 µL of TransfeX (ATCC, ACS-4005) was added to the cocktail, and the mixture was left to sit for ∼15 minutes. Following this period, the cocktail was added to the 1 mL of isolated HEK293T cells, well-mixed, and plated onto poly-D-lysine (PDL) or collagen-coated cell culture dishes until ready for experiments. Transfected cells were then used for calcium imaging experiments 48-120 hours post-transfection. Successful transfection was verified by GFP expression. Non-transfected cells were plated on to poly-d-lysine (PDL) or collagen-coated coverslips immediately following trypsinization. All cells prepared for imaging were maintained in 1 mL of the same phenol red-free DMEM used to grow them.

### Retina isolation

All animal procedures were performed under an Institutional Animal Care and Use Committee (IACUC)-approved protocol at the University of Illinois Urbana-Champaign. Mice with eGFP-expressing ipRGCs (Prof. Tiffany Schmidt, Northwestern)^27,28^ on a hybrid B6/S129S background were used for these investigations. Opn4-GFP^+^ mice were bred with F1 B6/S129S mice (The Jackson Laboratory, Strain #: 101043). Tail clippings were collected from generated litters after weaning (P21) and outsourced for genotyping (TransnetXY). Opn4-GFP-expressing mice aged above P21 were anesthetized following 1-3% isoflurane induction and euthanized by cervical dislocation. Eyes were immediately enucleated and placed in sterile-filtered Ames Medium (Osmolarity ∼300-310) for retina isolation. The retina was removed using established procedures^76,77^. The cornea was first punctured to reduce intraocular pressure, and then removed with fine scissors. The lens was then removed from the eye cup, along with the optic nerve and all attached muscles. The eye cup was then placed in an enzyme solution containing 5 mg/ml protease (Sigma, P4032), 1 mM L-Cysteine (Sigma), and 0.5 mM EDTA (Sigma) in Eagle’s Modified Essential Medium (EMEM) for 10-20 minutes. The tissue was then transferred to an inhibitor solution containing 2 mg/ml ovomucoid and 10% FBS in EMEM for 10 minutes. The eye cups were then rinsed 3x in Ames Medium and transferred to a separate container containing carbogenated Ames Media. The retinae were then isolated and further processed for calcium imaging experiments. Ames Medium was continuously perfused with carbogen (95% O_2_, 5% CO_2_).

### Calcium imaging

For HEK293T cell cultures, a final concentration of 5 µM of the red-shifted calcium indicator Calbryte-590^47^ (dissolved in DMSO) was loaded to each individual imaging dish, along with 0.05% Pluronic F-127. The indicator was maintained in the cell culture media for 1 hour. Thereafter, the cell culture media was removed, and 1 mL of fresh DMEM was added to each culture. To ascertain if sufficient retinal tissue was present for melanopsin activation, 20 µM *9-cis* retinal was added to the cell culture media for at least 30 minutes prior to imaging. Average power for calcium imaging (1030 nm laser source) was maintained at ∼25 mW, unless otherwise reported.

Calcium loading was performed in retina following isolation from the eye cup, similar to protocols in previous literature^76^. For each individual retina present, 20 µM Calbryte-590 and 0.20% Pluronic F-127 was loaded for 2 hours while aerated with carbogen (without bubbling) to ascertain widespread presence in all RGCs. For a pair of retinas, this equates to 40 µM Calbryte-590 and 0.40% Pluronic F-127 for incubation. After 2 hours, the retinae were rinsed three times in Ames Media and stored in carbogenated Ames Media until the imaging sessions. Retina slices were then placed under the microscope for image capture and stimulation. During imaging, retinae were incubated with ∼4 µM 9-cis retinal. Retinae were also incubated with a cocktail of synaptic inhibitors, unless stated otherwise, to prevent input from rods and cones^59^, although absorption is unlikely due to the high axial confinement of two-photon absorption^38^. The cocktail consisted of ∼50 µM D-2-amino-5-phosphonovalerate (D-AP5, Tocris), ∼20 µM 6,7-Dinitroquinoxaline-2,3-dione (DNQX, Tocris), and ∼8 µM Di-hydro-β-erythroidine (DhβE, Tocris). Average power was maintained at ∼50 mW, unless otherwise reported. This was optimized ensuring minimal to no bleaching of melanopsin (Supplementary Fig. 13), maximization of signal-to-noise ratio (Supplementary Fig. 14), and clear visualization of calcium transients resulting from ipRGCs (Supplementary Fig. 15).

### Data analysis

All imaging data was analyzed using custom MATLAB scripts. Calcium imaging transients were analyzed using a previously-developed algorithm^78^. Briefly, a 5-pixel diameter circular region from cell bodies of interest was used to generate calcium transients. The mean pixel intensities from these regions were calculated for each frame and assessed across each set of video frames to generate a trace of calcium activity over time. Normalization was calculated as 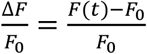, where F(t) denotes the mean fluorescence intensity at a given time, and F_0_ represents the baseline fluorescence signal from a given cell. This quantifies a percent-change in baseline fluorescence following imaging onset. The process was repeated for all cells of interest. A biexponential fit of the form *A*(*e*^−*bt*^ − *e*^−*ct*^) was applied to the calcium imaging data from the HEK293T cells, and (1 − *a*)*e*^−*bt*^ + *a* for isolated retinae, after normalization to the 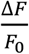 maximum to fit for activation and deactivation kinetics. The peak magnitude of the response following 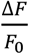 quantification was also determined. Biexponential fits, as reported in the main text, were applied to averages of multiple calcium traces for a given experimental condition. That is, for all transients from a given power or scan duration, all stimulation epochs for this condition were averaged and a fit was applied to that average. This was done to produce a more faithful quantification of the true rise and decay times for all transients, which otherwise show wide variability with biexponential fitting (Supplementary Figs. 8-10) due largely to the persistent response of melanopsin in HEK293 cells. Biexponential fits applied to individual calcium epochs are included to demonstrate this variability (Supplementary Figs. 8-10), but are not used for quantifying and interpreting activation kinetics, which are otherwise performed on the average trace for each experimental condition (Figs. 1 and 2). Statistical analysis was performed in MATLAB. A two-tailed Student’s t-test was used to quantify statistical differences in kinetics information.

For visualization of the supplementary videos, the recordings were denoised using a self-supervised deep learning framework (i.e., DeepCAD-RT)^79^. By exploiting the spatiotemporal redundancy within calcium imaging signals, the DeepCAD-RT suppresses detection noise and improves the signal-to-noise ratio (SNR) of calcium recordings without requiring high-SNR ground truth for training. In this study, each calcium recording was divided into two sub-stacks (i.e., odd-numbered frames and even-numbered frames), which were then used as the input volume and the target volume to train the deep neural network. The backbone of DeepCAD-RT is a simplified 3D-UNet^80^, of which the features were pruned for higher processing speed and reduced memory consumption. The training and inference procedures implemented in this study mainly followed the original DeepCAD-RT configurations, except for the loss function, the padding mode, and the inference order. The loss function was changed from the arithmetic average of L1- and L2-norm loss to only L2-norm loss for enhanced reliability of model outputs under photon-starving conditions. The padding mode of convolutional layers inside the 3D-UNet was changed to reflect-padding, eliminating edge artifacts caused by zero-padding in the restored calcium images. During model inferencing, a two-step inference procedure was implemented, which generated even-numbered and odd-numbered frames separately. The two restored sub-stacks were then combined to form the final restored calcium recording.

## Data availability

The data that support these results are available from the corresponding author upon reasonable request and through collaborative investigations.

## Code availability

The code used to analyze the data in this work is available from the corresponding author upon reasonable request and through collaborative investigations.

## Acknowledgements

The research reported in this publication was supported by training grants from the National Institute of Biomedical Imaging and Bioengineering (NIBIB) and the National Institutes of Environmental Health Sciences (NIEHS) of the NIH under Award Numbers T32EB019944 and T32ES007326. The content is solely the responsibility of the authors and does not necessarily represent the official views of the National Institutes of Health. This research was also supported in part by grants from the Air Force Office of Scientific Research (FA9550-17-1-0387) and the NIH/NIBIB Center for Label-free Imaging and Multiscale Biophotonics (CLIMB) (P41EB031772), including a Supplement grant. Additional information can be found at http://biophotonics.illinois.edu.

## Author information

### Contributions

C.A.R. and S.A.B. conceived and directed the study. C.A.R., R.R.I., and Y.Z.L. developed the optical imaging and control system used in this study. C.A.R., B.T., and J.K. performed cell culture and transfection experiments. C.A.R. performed murine retina experiments. C.A.R. prepared samples for imaging and performed imaging experiments. C.A.R. and J.S. processed and analyzed the imaging data. C.A.R., E.J.C., and E.A. performed murine husbandry, breeding, and genotyping of the animals used in this study. H.A. generated and prepared the plasmid used in this study. H.A. and P.R. provided the resources for plasmid use in this study. T.S. generated and donated the mouse lines used in this study. C.A.R. prepared the original draft and figures in the manuscript. All co-authors assisted in providing feedback on the manuscript. S.A.B. provided financial support and the resources used to pursue all experiments in this study.

## Ethics declarations

### Competing interests

The authors declare that they have no competing financial interests.

## Notes

### Competing Interest Statement

The authors have declared no competing interest.

