## Supplementary information for "Two-photon activation, deactivation, and coherent control of melanopsin in live cells"

[1]Beckman Institute for Advanced Science and Technology
[2]Department of Bioengineering
[3]Department of Engineering Physics
[4]Department of Molecular and Cellular Biology
[5]Department of Electrical and Computer Engineering
[6]Neuroscience Program
[7]NIH/NIBIB P41 Center for Label-free Imaging and Multiscale Biophotonics (CLIMB)
[8]University of Illinois Urbana-Champaign, Urbana, IL
[9]University of Maryland, Baltimore County, Baltimore, MD
[10]Northwestern University, Evanston, IL

**Supplementary Note 1:**

*Melanopsin photocycle:*

Among the unique features of melanopsin, one of the most compelling that has recently been brought to light is its photoswitchability. That is, melanopsin can be excited with certain wavelengths of light (notably blue light), and deactivated with others (yellow to red)[30–32]. Notably, it has also been proposed that melanopsin exists in three states to permit these transitions. It begins in a resting state that is only achievable through incubation in prolonged darkness[30,33], which is typically referred to as the $|R\rangle$ state. Following absorption of a high-energy photon, namely blue light, it is activated, entering the metamelanopsin state, or $|M\rangle$ state. Again, melanopsin can revert to the $|R\rangle$ state from the $|M\rangle$ state following a prolonged absence of light exposure. Alternatively, melanopsin can be transitioned to a second silent state through exposure to red-shifted light, (at or above 561 nm). This final state is known as the $|E\rangle$ state, or extramelanopsin state, although its existence in a warm biological environment is yet to be identified[30,33]. Whether or not this state is the true state is still up for debate, namely due to the proposed structure of the chromophore. That is, the $|R\rangle$ state begins with the *11-cis* retinal isomer. Following activation to the $|M\rangle$ state, it transitions to *all-trans* retinal. Retinal in the $|E\rangle$ state has been proposed to be in the *7-cis* retinal state, which has only been achieved experimentally in highly-controlled experimental conditions[33], making the existence of retinal in this state following exposure to 561 nm light highly debated. The $|E\rangle$ is also theorized to naturally relax to the $|R\rangle$ state. Nonetheless, there is strong evidence for the tri-stability of melanopsin independent of the exact retinal isomer[30,34,35] from multiple research groups to date, in different cell types. A diagram of this photocycle and the retinal isomers associated with each state is illustrated in Supplementary Fig. 1A, along with the proposed two-photon control pathway detailed in this work (Supplementary Fig. 1B).

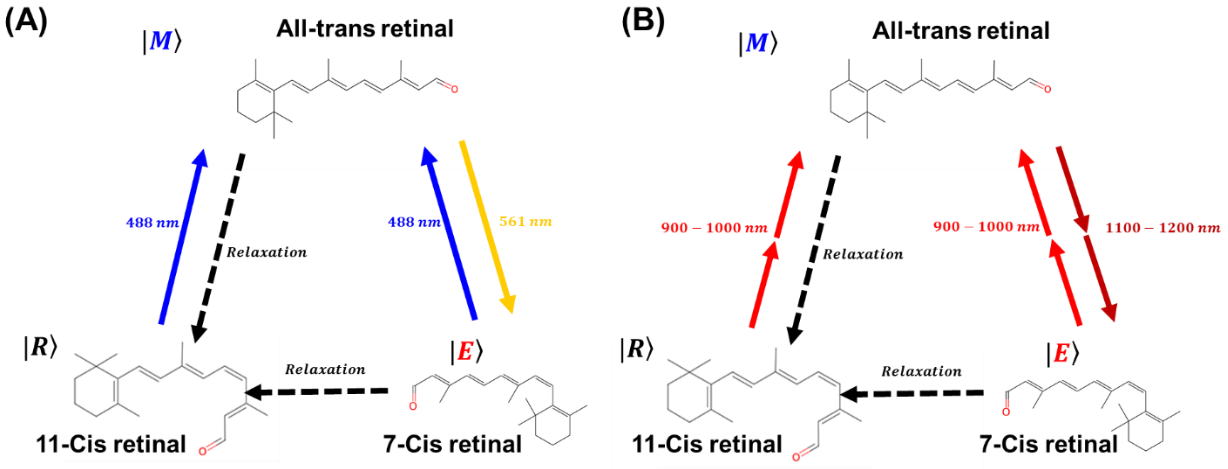


**Supplementary Fig. 1: Melanopsin photocycle illustration. a,** In the visible light regime, the retinal chromophore begins in the 11-cis retinal state, which isomerizes to the all-trans state following absorption of a photon in the visible regime. This is maximally absorbed at ~488 nm, as illustrated in the diagram. In this state, the all-trans retinal can absorb a lower energy photon, maximally at 561 nm, to drive it to a third, inactive state, currently theorized to be the 7-cis retinal isomer. The chromophore can be dynamically transitioned between these two states using higher and lower energy photons, and driven back to the baseline ground state of 11-cis retinal following incubation in prolonged darkness. **b,** The same process, in theory, can be elicited in the two-photon regime using infrared photons that bridge this energy gap, and using the principle of coherent control described in this work.

**Supplementary Note 2:**

*Multiphoton absorption and coherent control theory:*

The principal phenomena leveraged in this work are two-photon absorption and coherent control of two-photon transitions for control of photochemical reactions. Two-photon absorption is a phenomena by which two photons of energies equal to roughly half the energy of electronic transitions in a molecule interact to be absorbed in a chromophore[1–3]. These and other multiphoton transitions can be predictably controlled through a process known as coherent control. Coherent control, broadly, is a process that involves shaping the spectral phase of a light source to control a photochemical reaction. That is, light can be manipulated to modulate the efficiency of a photoproduct, its yield, and potentially the photoproduct itself[4–6]. The theory was formally described in the spectral and temporal domains[7–9] by Shapiro & Brumer and Tanner & Rice, respectively. Coherent control has found applications in quantum computing[10,11], spectroscopy[12,13], and microscopy[14–20] among other fields. Most relevant in this work is how it can be used to control the photoreaction of multiphoton processes interacting with a chromophore, described and demonstrated originally by Meshulach and Silberberg in cesium[21]. This model was expanded upon through multiphoton intrapulse interference[22–24]. A mathematical model can be developed by conceiving this nonlinear interaction as two NIR photons interacting with each other (simultaneous absorption) while interacting with a nonlinear medium (the two-photon chromophore). The initial interaction between the two photons, using an impulse-response model, can be treated as a convolution between the two photons. That is:

$$E^{(2)}(2\omega_0) \approx \int_{-\infty}^{\infty} E(\omega_0 - \Omega)E(\omega_0 + \Omega)d\Omega = E(\omega) * E(\omega).$$

Using the convention adopted by Lozovoy *et. al.*[22], where $\Omega$ denotes spectral detuning from the central wavelength of our laser, $\omega_0$, such that $\Omega = \omega - \omega_0$. Here, * denotes the convolution operator for convenience and readability. In the case of coherent control of two-photon interactions, the interaction between two photons of a given wavelength and a chromophore with two-photon cross-section $\sigma_2$ can be modelled as a convolution of their spectra and spectral-dependence:

$$S^{(2)} \approx \int_{-\infty}^{\infty} \sigma_2(\Delta)|E^{(2)}(\Delta)|^2 d\Delta = \sigma_2(2\omega) * E^{(2)}(2\omega)$$

where $\Delta = \omega - 2\omega_0$ describes the electronic transition invoked in the case of two-photons. Consider here that the electric field $E(\omega)$ is complex-valued, as the phase becomes less arbitrary when individual components are modulated with respect to the central wavelength of your pulse. Assuming $E(\omega) = Ae^{i\varphi(\omega)}$ where $A$ is the amplitude of the pulse, and $e^i$ denotes the complex exponential, this expands the above relationship to:

$$S^{(2)} \approx \int_{-\infty}^{\infty} \sigma_2(\omega - 2\omega_0)|A|^2 e^{i[\varphi(\omega+2\omega_0)+\varphi(\omega-2\omega_0)]} d\omega$$

$$= \sigma_2(2\omega) * E^{(2)}(2\omega)e^{i[\varphi(\omega+2\omega_0)+\varphi(\omega-2\omega_0)]}.$$

The phase-dependence of the two-photon signal implicates that the signal resulting from a nonlinear interaction including two or more photons can be modulated through tuning of the spectral phase of the light source. This makes the technique generalizable for any chromophore or harmonophore involved in a multiphoton interaction.

To determine if tailoring the phase profile can cause changes in melanopsin activation, it is important to understand the underlying photophysics. To this end, we modelled the two-photon interaction by modelling second-harmonic generation (SHG), using a Dirac delta. In our model, this is represented with a value of 1 centered at the central wavelength of the source. The measured spectrum from our supercontinuum was used to model the spectrum of the electromagnetic field in the frequency domain used for simulations. The effects of different dispersion profiles on SHG signals are shown in Supplementary Fig. 2. Of special interest is GDD, shown experimentally in Supplementary Fig. 3 (C-F), and third-order dispersion (TOD) on the temporal pulse duration and shape. This is shown in Supplementary Fig. 1 (left), and second-order signal, shown in Supplementary Fig. 1 (right) for different TOD. With GDD, the temporal duration of an ultrafast pulse increases in proportion to the amount of GDD imposed on the pulse. This, in effect, manifests itself into lower SHG and two-photon absorption signals with greater degrees of GDD. For SHG and two-photon absorption signals, this trend continues with TOD, where higher degrees of TOD cause drastic drops in SHG and two-photon signals. In the temporal domain, pulse duration is also affected, but rather than a uniform dispersion of wavelengths across the duration of the pulse, all wavelengths beyond that of the central wavelength disperse in the same temporal direction according to the degree of TOD. Intuitively, this can be reasoned to be a result of the linear region of the profile-imposed phase on our spectrum. That is, wavelengths above or below the central wavelength of our pulse are imposed with the exact same slope/gradient (that is, the rate of change) about the central wavelength. For positive TOD, they receive the same positive slope and are delayed, whereas for negative TOD, the slope is negative, and the wavelengths are advanced about the central wavelength. In contrast, pulses imposed with GDD receive the same degree of phase gradients of different signs. For positive GDD, wavelengths below the central wavelength receive a negative slope, and those above receive a positive slope. This reverses in the case of a negative slope, causing wavelengths to delay or advance in opposite directions.

The SHG and two-photon signals can be explained through multiphoton intrapulse interference[22], where the pulses interfere with the nonlinear media (BBO crystal or chromophore for SHG and two photon absorption respectively) in such a way that they interfere and can cause predictable changes in these media. This interaction becomes especially interesting for non-centrosymmetric phase profiles, such as TOD or phase-shifted cosinusoidal phase profiles among others [17,22,23,25–27], illustrated in Supplementary Fig. 1 (A-C, right) for TOD. By keeping the degree of dispersion fixed, and tuning the central wavelength in the case of TOD, the pulses interfere in a nonlinear medium such that signal is maximized about the central wavelength, $\omega_0$, that the TOD is tuned to, according to the third-order dispersion term $a_3(\omega - \omega_0)^3$. In this simulation, we maintained $a_3 = 15{,}000\ fs^3$. For cosinusoidal shapes that take the form $\alpha\cos\,(\gamma t - \delta)$, a similar phenomenon arises such that the anticipated peak excitation wavelength is tuned by tunning the $\delta$ term. These profiles provide researchers with a unique and simple physical tool to predictably control any photochemical interaction in the multiphoton regime. This has yet to be demonstrated for predictable control of opsin kinetics, so we applied this tool to determine if this biophysical interaction which has been demonstrated in fluorescent chromophores[17,22,23,25–28] translates to these more complex, non-radiative systems.

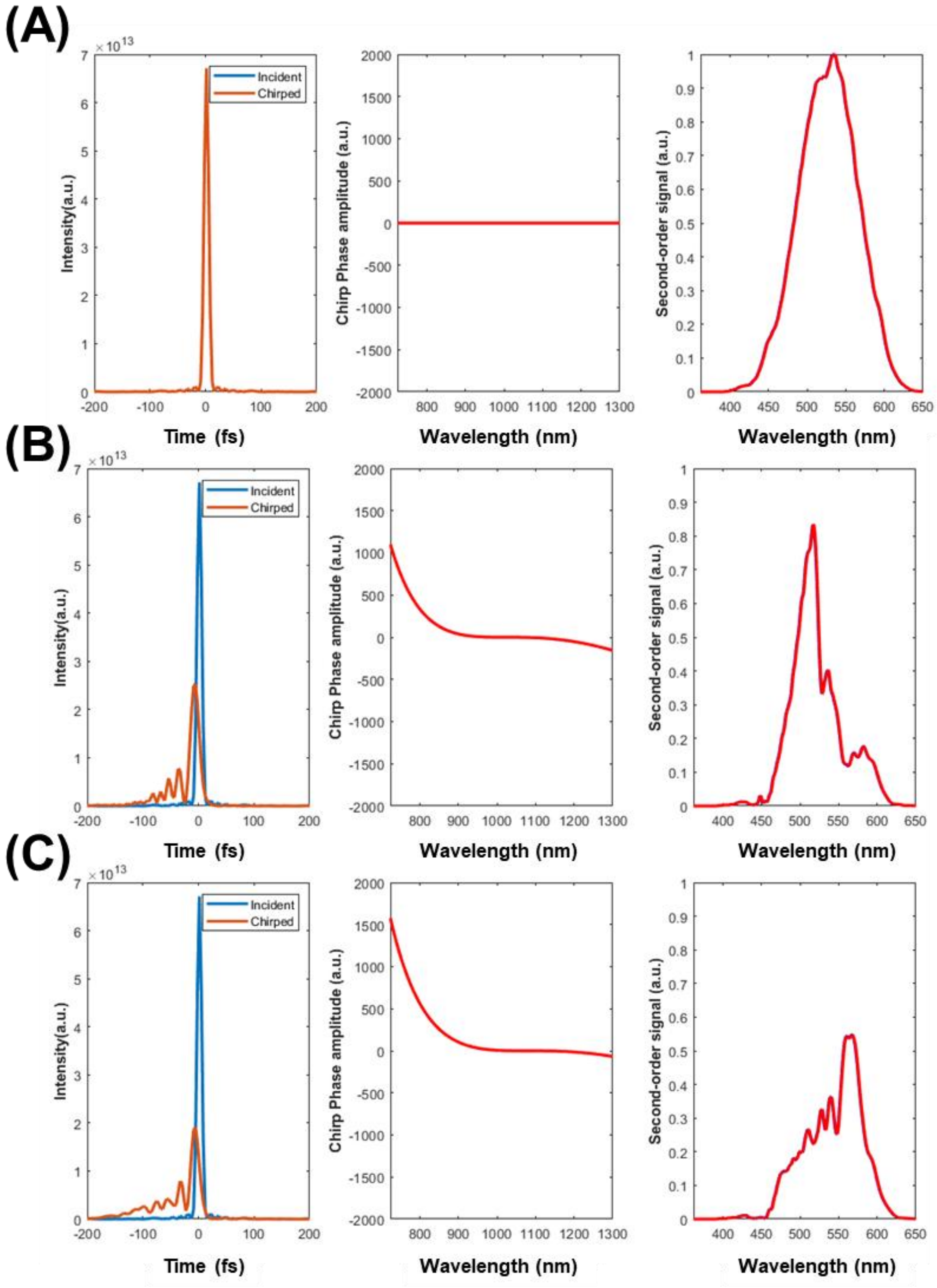


**Supplementary Fig. 2: Effect of phase profiles on two-photon signals. a,** transform-limited signal **a** results in temporal profiles and maximum two-photon signal intensity (left). This is denoted by a flat phase profile (center), with a second-order signal that spans the supercontinuum spectrum (right). **b,** Following the application of TOD, the temporal profile changes such that all photons are delayed in the same temporal direction in time (left) when centered at the center wavelength (middle). The signal results in peak signals still centered around the central wavelength of the source (right), but reduced efficiency in other spectral windows (right). **c,** When TOD is centered at a longer wavelength (middle) the temporal profile does not change much (left), but the second-order signal changes such that the peak signal is shifted towards the new central wavelength (right), while other contributing wavelengths are reduced.

Ascertaining that the pulse is at its near-transform limit (TL) is critical for proper two-photon coherent control of a chromophore. Here, our supercontinuum pulses were compressed to their near transform-limit using the multiphoton intrapulse interference phase scan (MIIPS) algorithm[28], and verified using frequency-resolved optical gating (FROG)[29]. Results from running MIIPS on the supercontinuum from our PCFs (Supplementary Fig. 2B) along with the corresponding frequency-resolved optical gating (FROG) trace (Supplementary Fig. 2C) and autocorrelation (Supplementary Fig. 2D) are shown in Supplementary Fig. 2 showing a final pulse width of ~20 fs full-width half-maximum (FWHM), near the transform limit for this supercontinuum pulse.

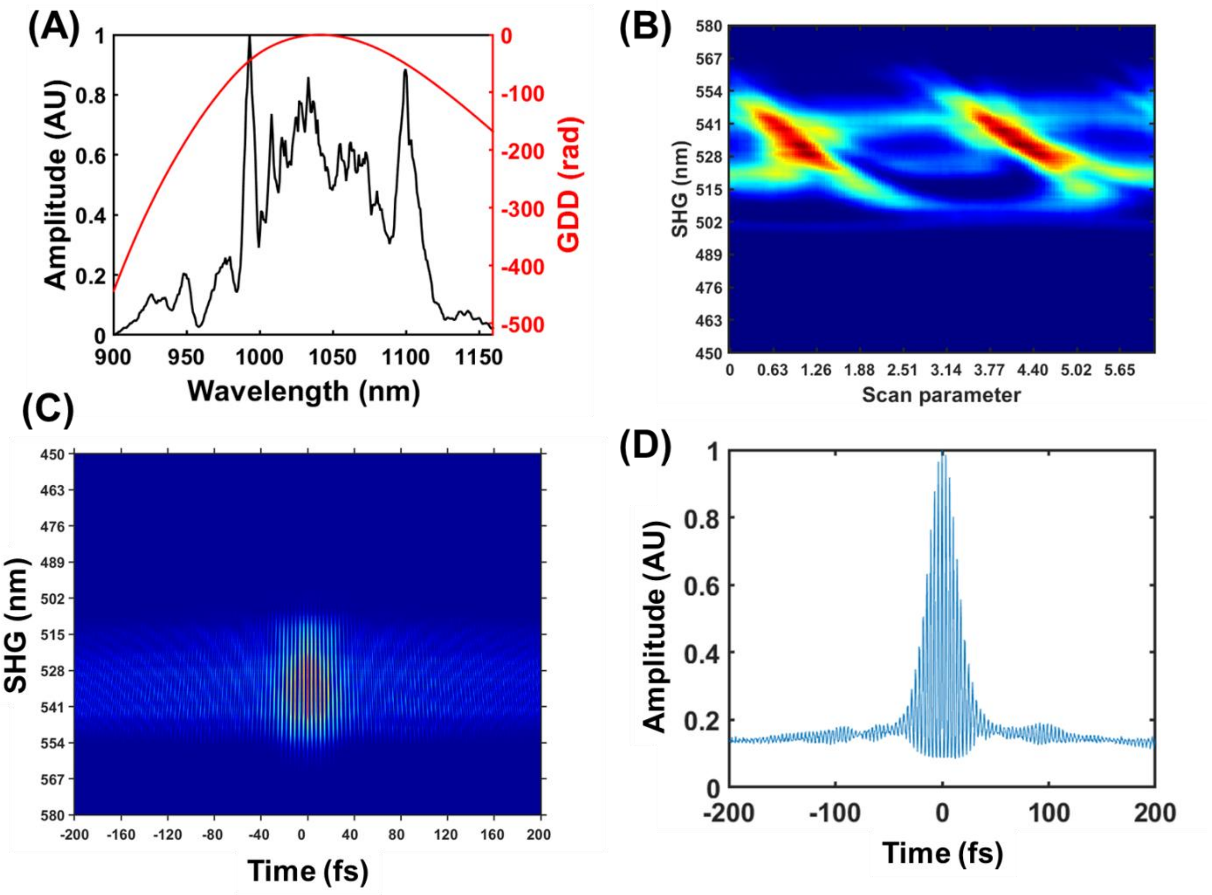


**Supplementary Fig. 3: Supercontinuum pulse characterization. a,** Measured supercontinuum used in this study (black) and the spectral phase measured following MIIPS protocol to generate TL pulses (red). **b,** MIIPS trace measured following pulse compression. The diagonal trace is characteristic of TL pulses. **c,** FROG trace measured after pulse compression. **d,** The corresponding autocorrelation function for **c**, integrated across all wavelengths, with a full-width half maximum of ~20 fs in duration.

To ensure that phase modulation using this experimental system could induce predictable changes in nonlinear signals, we tested the effects of varying degrees of GDD on SHG in a Barium Borate (BBO) crystal, illustrated in Supplementary Fig. 3 The full supercontinuum spectrum and measured spectral phase to compensation for dispersion are shown in Supplementary Fig. 3A. Supplementary Fig. 3B shows the temporal duration of ~20.3 fs for the pulse following dispersion compensation. Following MIIPS to compress these optical pulses, results in Supplementary Fig. 3C show an increase in the SHG signal compared to baseline, demonstrating success of MIIPS for pulse compression but also a fundamental form of coherent control. The relative difference between uncompensated and compensated SHG signal intensity is shown in Supplementary Fig. 3D. By adding varying degrees of GDD, we also see symmetric changes in the SHG for positive and negative chirp in Supplementary Fig. 3E, an expected result and a demonstration of predictable control of nonlinear signals by phase modulation. Lastly, using a cosinusoidal phase profile, such as that used to run MIIPS for pulse compression, we see in changes in Supplementary Fig. 3F in the maximum and minima SHG signal that track with simulated responses, further demonstrating predictable control of nonlinear effects even with more complex phase functions. These collectively demonstrate the reliability of this system, illustrated in Supplementary Fig. 4, for experiments in more complex systems, and establish high-quality system performance for the coherent control of melanopsin experiments performed in this study.

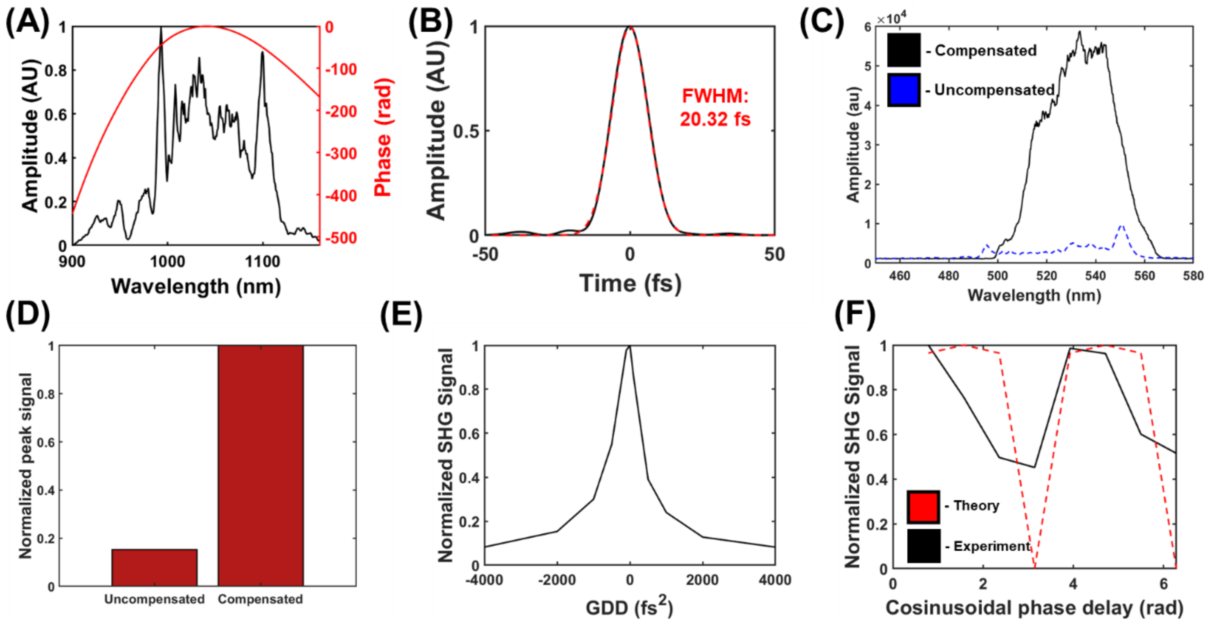


**Supplementary Fig. 4: Testing effects of phase modulation on SHG signal from BBO crystal.** **a,** Supercontinuum spectrum and phase applied to spectrum. **b,** Calculated pulse-width following MIIPS compression. **c,** SHG spectrum obtained before (blue) and after (black) dispersion compensation with MIIPS. **d,** Normalized bar-graph showing the difference in signal between the uncompensated and compensated pulses. **e,** Effects of varying GDDs on SHG signal, normalized to the transform-limited signal at 0 GDD. **f,** Effects of manual cosinusoidal phase on SHG signal, akin to the profiles used for MIIPS characterization.

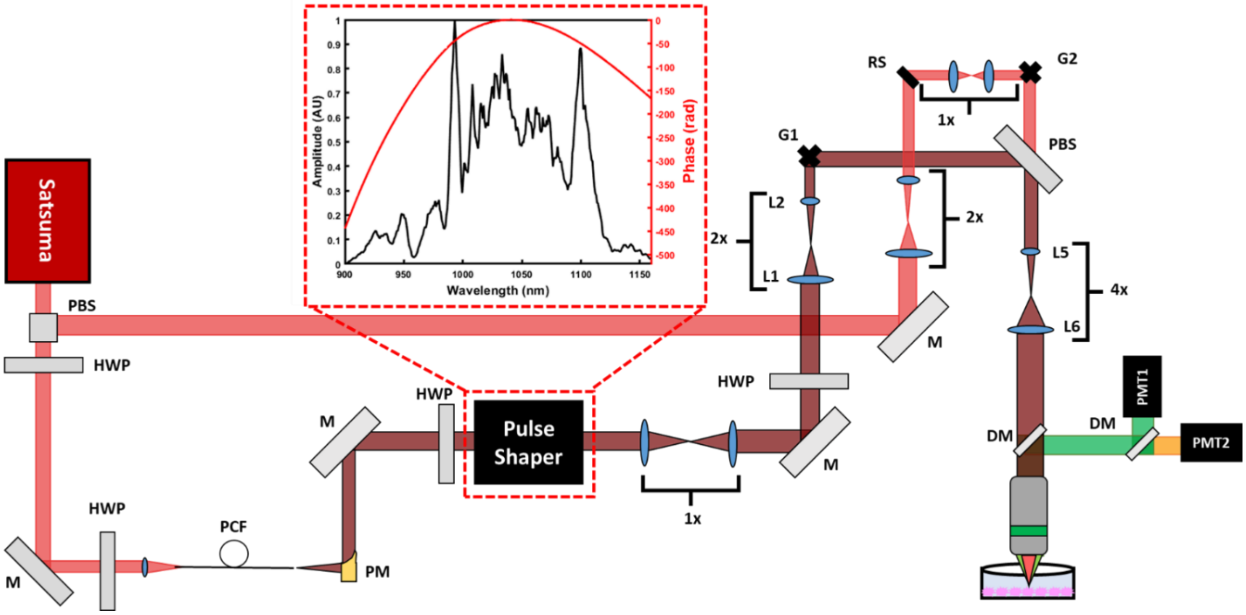


**Supplementary Fig. 5: System used for two-photon activation and imaging of melanopsin.** The laser source is split into two paths by a polarizing beam splitter (PBS). The first path is directed towards a resonant scanner (RS) and pair of galvanometers for rapid imaging. The other gets pumped to a PCF to generate a supercontinuum of light, which is redirected to the sample through a microscope objective for imaging. Both paths are used to generate images of calcium dynamics and GFP for the identification of melanopsin-expressing cells. Abbreviations: Polarizing beam splitter (PBS); Half-wave plate (HWP); Mirror (M); Photonic crystal fiber (PCF); Lens (L); Galvanometer (G); Resonant scanner (RS); Photomultiplier tube (PMT); Dichroic mirror (DM).

**Supplementary figures and data**

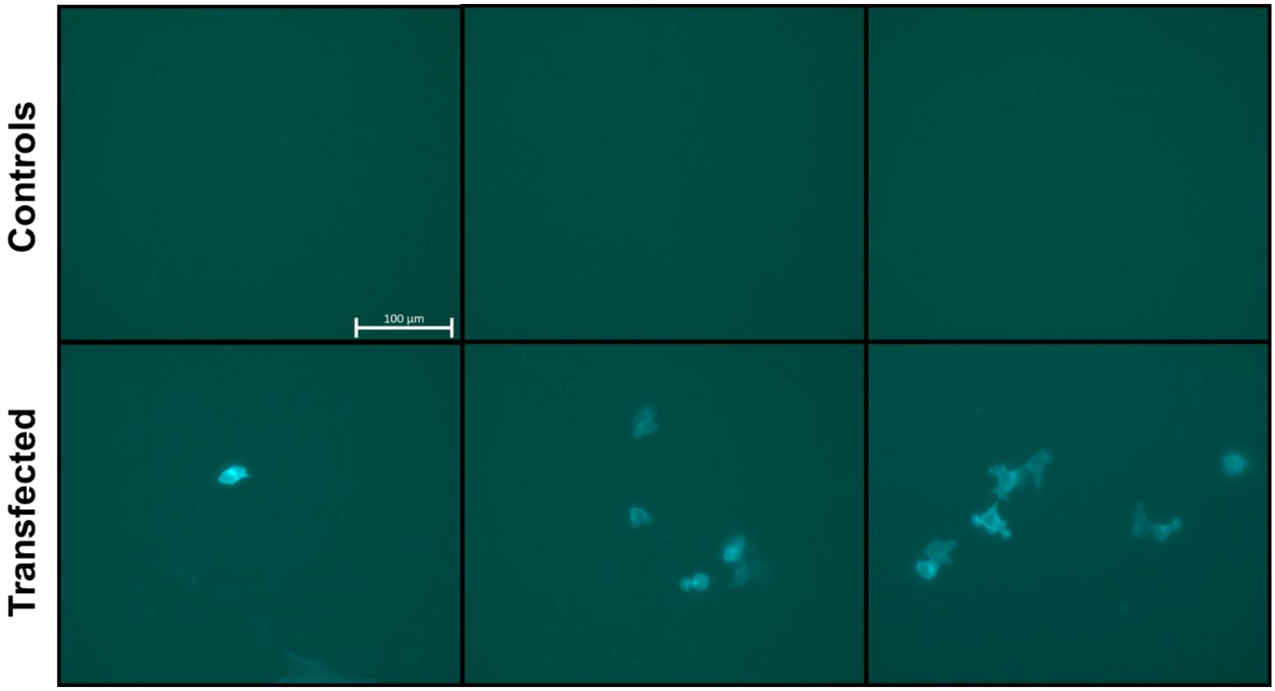


**Supplementary Fig. 6: Illustrating of melanopsin transfection.** The top panel shows controls, with no visible fluorescence at any field-of-view in the culture. The bottom panel shows a subset of HEK293T cells that do express GFP, and consequently melanopsin. Each column designates a different field-of-view for the same culture. Scale bar denotes 100 µm.

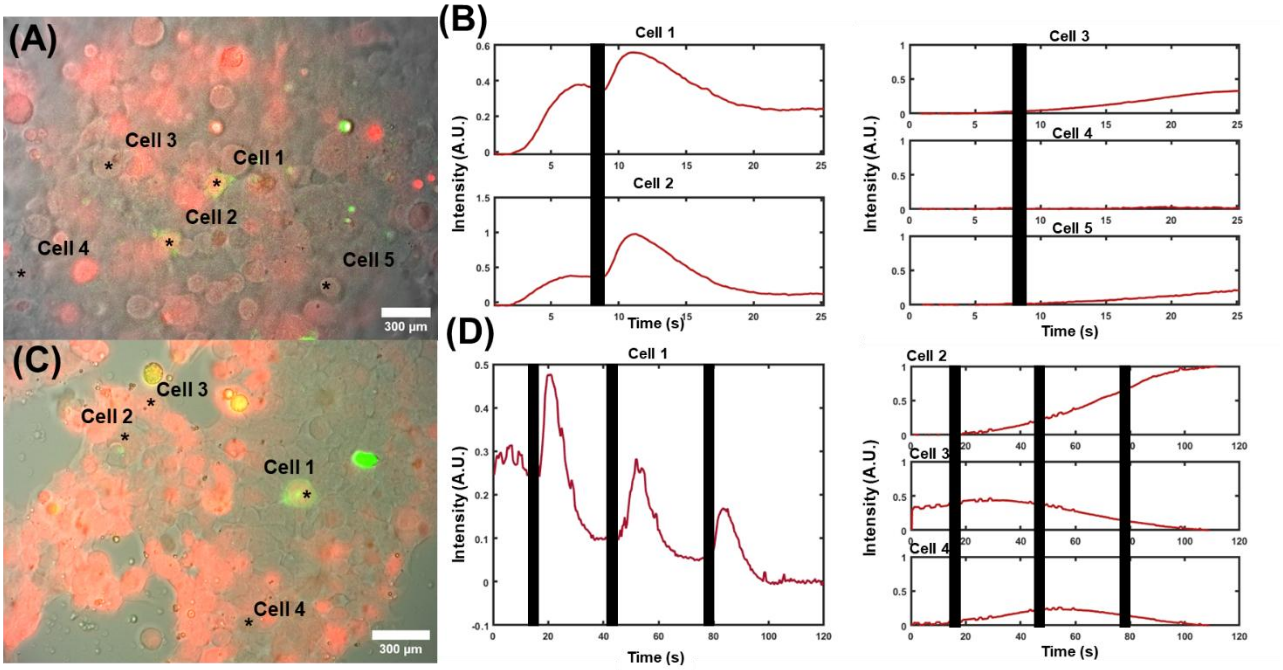


**Supplementary Fig. 7: Single-photon stimulation results. a,** Multimodal field-of-view of HEK293T cells with differential interference contrast (DIC) illumination, GFP fluorescence (green) indicating the presence of melanopsin, and the calcium indicator Calbryte-590 (red). **b,** Representative calcium transients from cells 1-5 in **a**. **c,** A different region of HEK293T cells expressing melanopsin. **d,** Evoked single-photon responses from melanopsin-expressing GFP cells, and adjacent controls. Black bars indicate instances when optical pulses were illuminating the cells. Results reported from n = 9 cells.

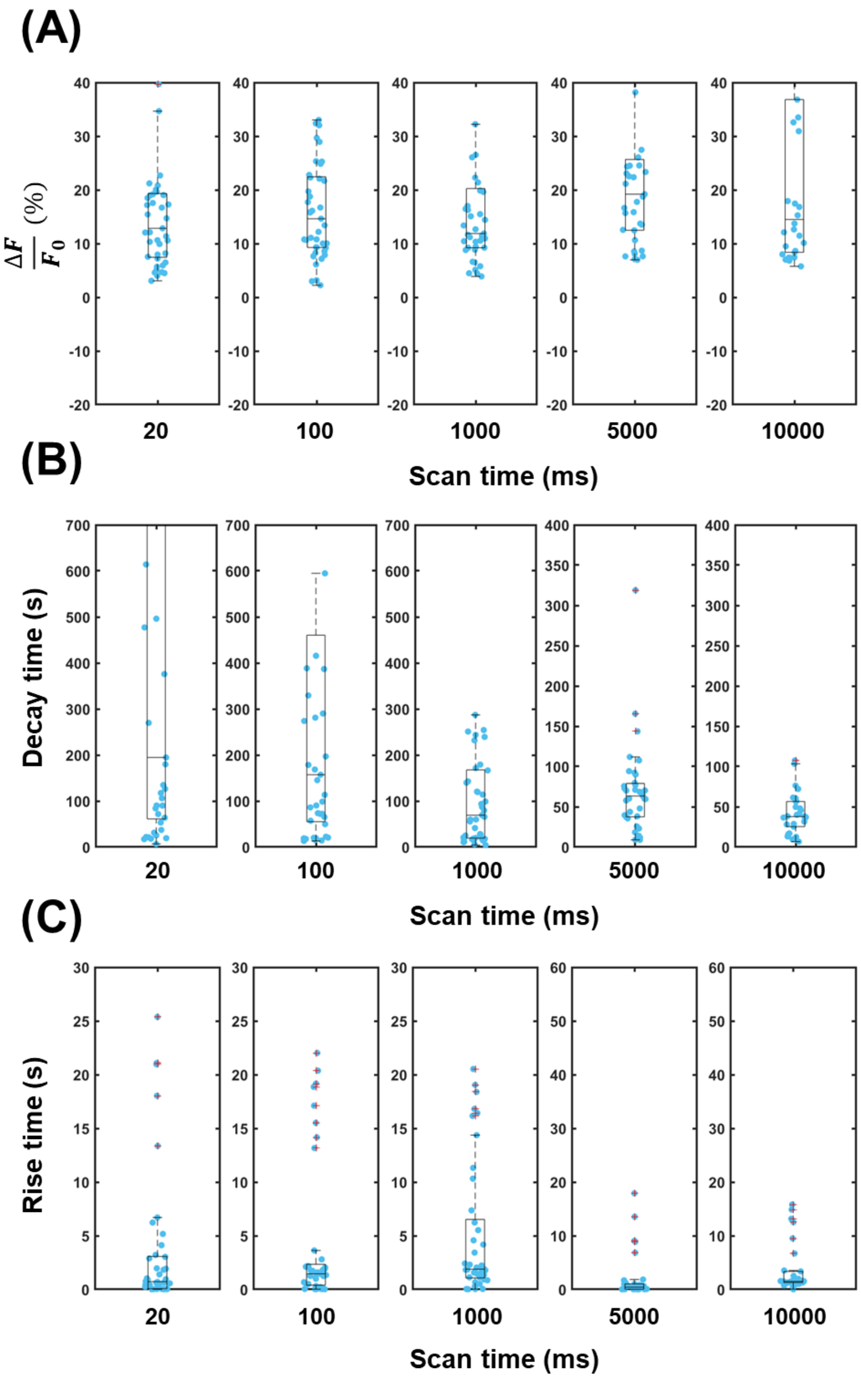


**Supplementary Fig. 8: Distributions of the decay times measured following exponential fitting of calcium imaging data following varying exposure times to the supercontinuum light. a,** The distribution of $\frac{\Delta F}{F_0}$ intensity for each spiral scan exposure duration. **b,** Decay time ($\tau_1$) of the calcium imaging traces for all individual traces. **c,** The distribution of calcium rise times ($\tau_2$) following varying exposure durations. The central line denotes the median, the bottom and top of the boxes the 25$^{th}$ and 75$^{th}$ percentile, respectively, and whiskers to non-outlier data extremes. Outliers are denoted by a red "+" symbol.

**Supplementary Table 1: Tabulated values for data reported in Fig. 1**.

| | **ΔF/F (%)**<br>**Mean** | **T1 (s)**<br>**Mean** | **T2 (s)**<br>**Mean** |
|---|---|---|---|
| **20 ms** | 7.62 | 71.6 | 1.6 |
| **100 ms** | 8.95 | 37.7 | 3.4 |
| **1000 ms** | 9.76 | 25.8 | 4.3 |
| **5000 ms** | 16.36 | 21.3 | 0.7 |
| **10000 ms** | 15.51 | 13.5 | 3.0 |

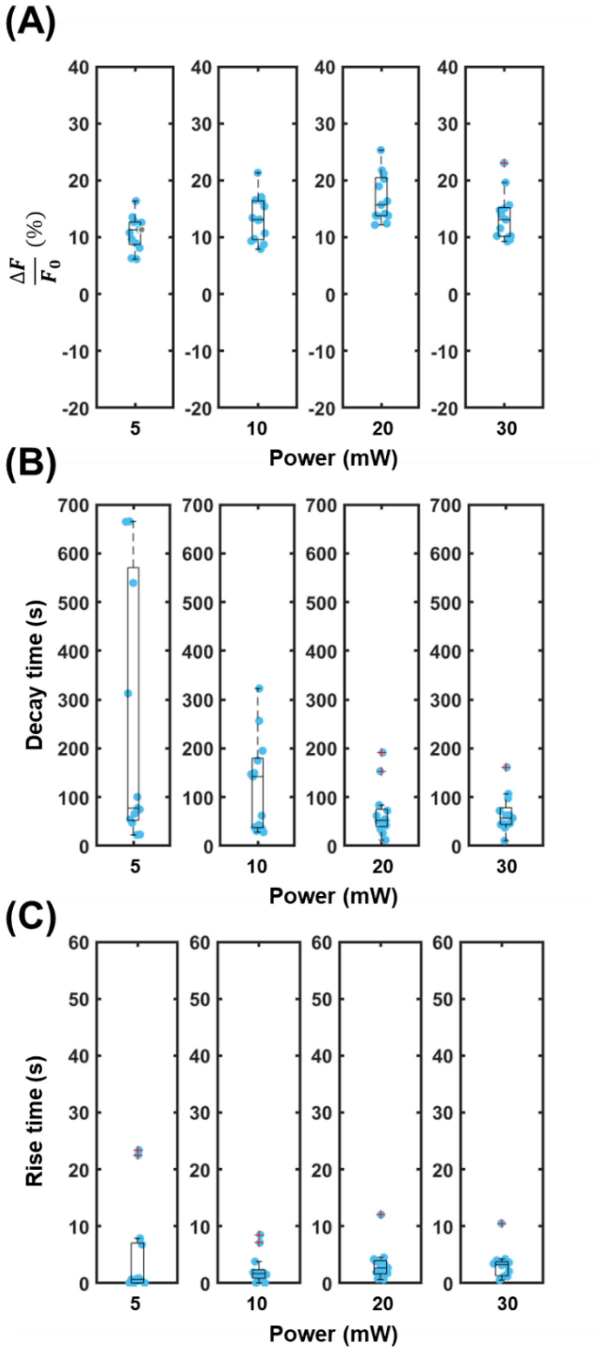


**Supplementary Fig. 9: Distributions of the decay times measured following exponential fitting of calcium imaging data at multiple incident powers. a,** The distribution of $\frac{\Delta F}{F_0}$ intensity for all individual traces following exposure to supercontinuum light at powers between 5-30 mW. **b,** Decay time ($\tau_1$) of the calcium imaging traces for all individual traces following exposure to supercontinuum light at powers between 5-30 mW. **c,** The distribution of calcium rise times ($\tau_2$) following exposure to supercontinuum light at different powers between 5-30 mW. The central line denotes the median, the bottom and top of the boxes the 25th and 75th percentile, respectively, and whiskers to non-outlier data extremes. Outliers are denoted by a red "+" symbol.

**Supplementary Table 2: Tabulated values for data reported in Fig. 2.**

| | **ΔF/F (%)**<br>**Mean** | **T1 (s)**<br>**Mean** | **T2 (s)**<br>**Mean** |
|---|---|---|---|
| **5 mW** | 5.78 | 54.0 | 1.4 |
| **10 mW** | 8.13 | 29.6 | 3.4 |
| **20 mW** | 11.10 | 25.3 | 3.4 |
| **30 mW** | 8.48 | 25.5 | 4.5 |

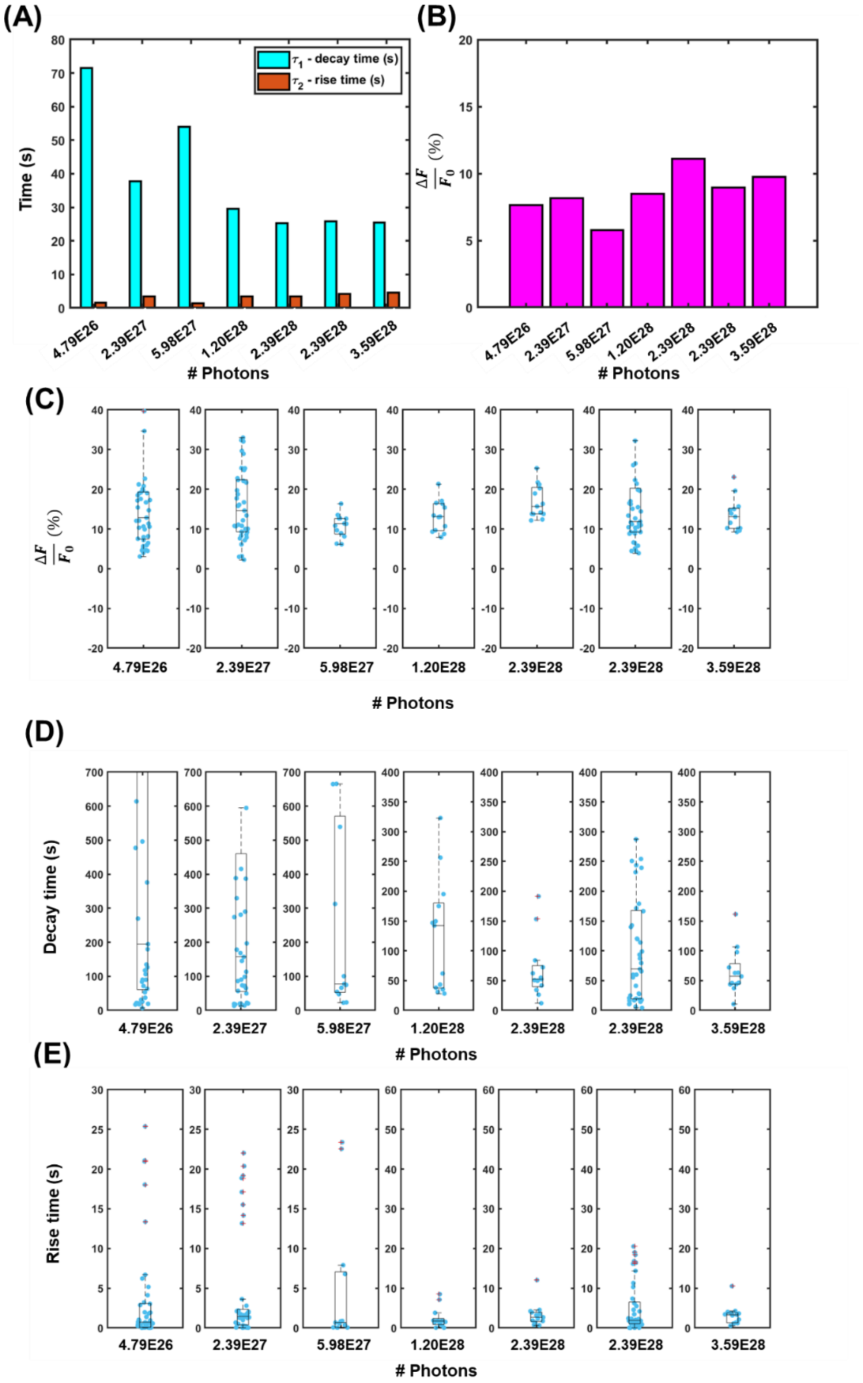


**Supplementary Fig. 10: Summary statistics of calcium imaging kinetics when separated by the number of irradiated photons at the sample. a,** The rise and decay times for calcium fitting following irradiation with varying incident numbers of photons. This was obtained from biexponential fits applied to a mean of all calcium imaging traces exposed to a given condition. **b,** Peak calcium transient for each incident number of photons. **c,** Distribution of peak calcium transient amplitudes for each condition of number of incident photons. **d,** Decay time ($\tau_1$) of the calcium imaging traces for all individual traces following exposure to supercontinuum light with different numbers of photons. **e,** The distribution of calcium rise times ($\tau_2$) following exposure to supercontinuum light with different numbers of photons.

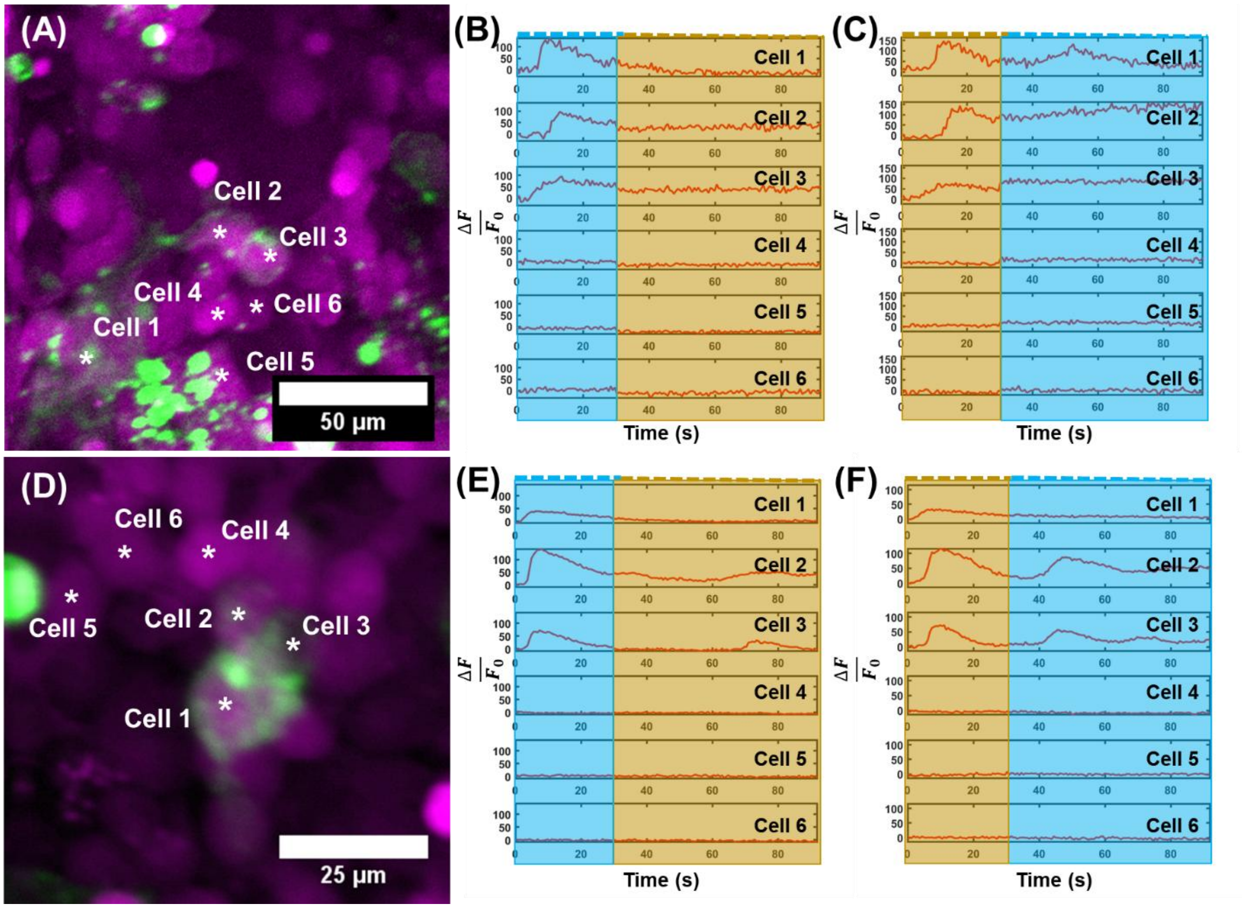


**Supplemental Fig. 11: Illustration of coherent control effects on melanopsin-expressing and non-expressing HEK293T cells. a,** A designated field-of-view. **b,** Transients from Cells 1-6 in **a** illuminated with TOD centered at 980 nm, then centered at 1140 nm after 30 s illumination. **c,** Illumination with supercontinuum in reverse order, initially with TOD centered at 1140 nm, then transitions to 980 nm. **d,** A designated field-of-view. **e,** Transients from Cells 1-6 in **d**, illuminated with a supercontinuum with spectral TOD centered at 980 nm, then 1100 nm. **f,** Reverse illumination paradigm, centered first at 1100 nm, then 980 nm. Cyan indicates illumination of supercontinuum anticipated for activation, and yellow/gold for deactivation kinetics. Responses evoked in n = 5 cells. Results reported from n = 12 cells.

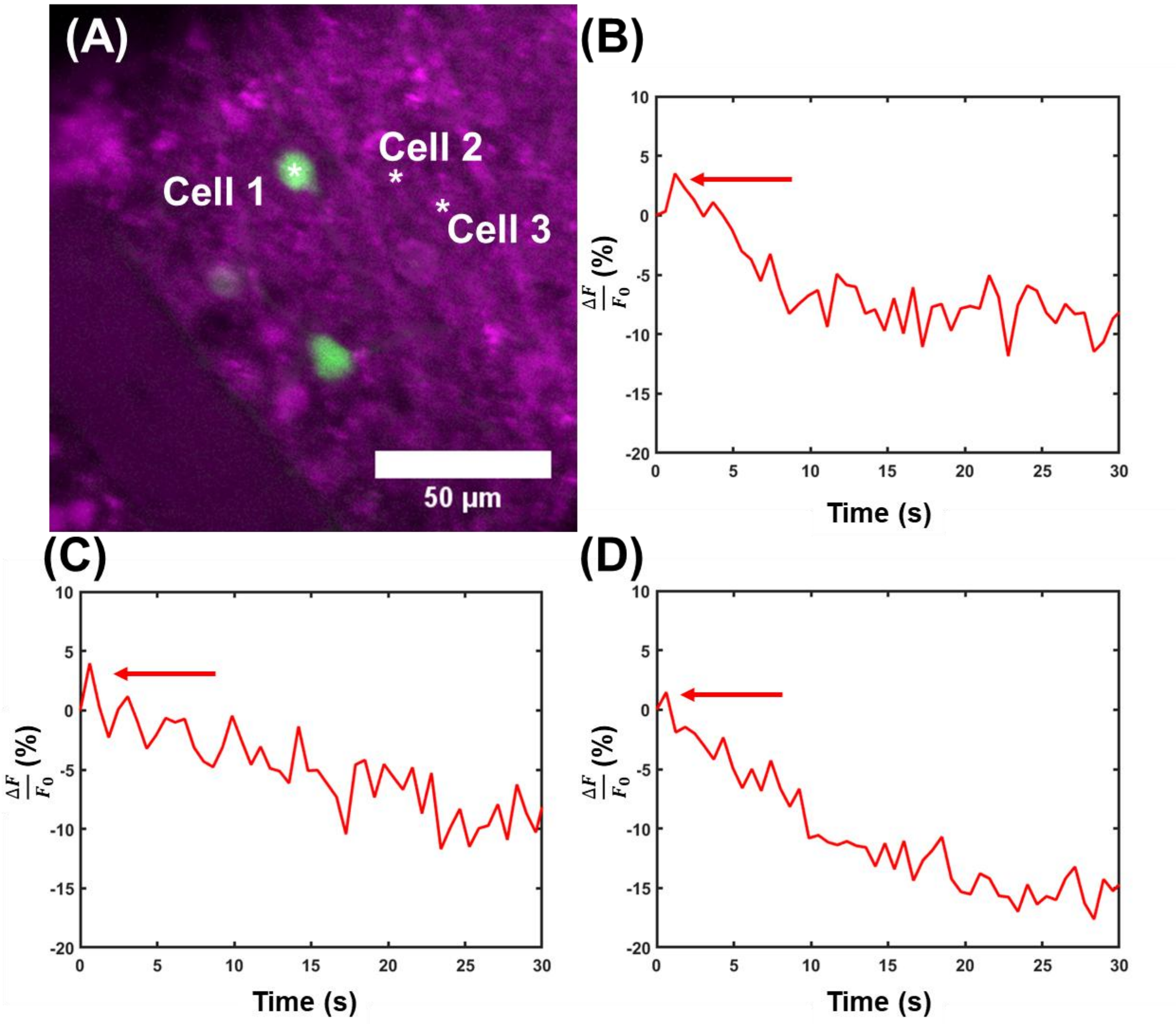


**Supplementary Fig. 12: Immediate rise in calcium transient for illuminated ipRGCs. a,** Retina region qualified for imaging. The resulting traces illustrate the immediate rise and subsequent decay of calcium levels for **b,** Cell 1, **c,** Cell 2, and **d,** Cell 3 for the retina region shown in **a**.

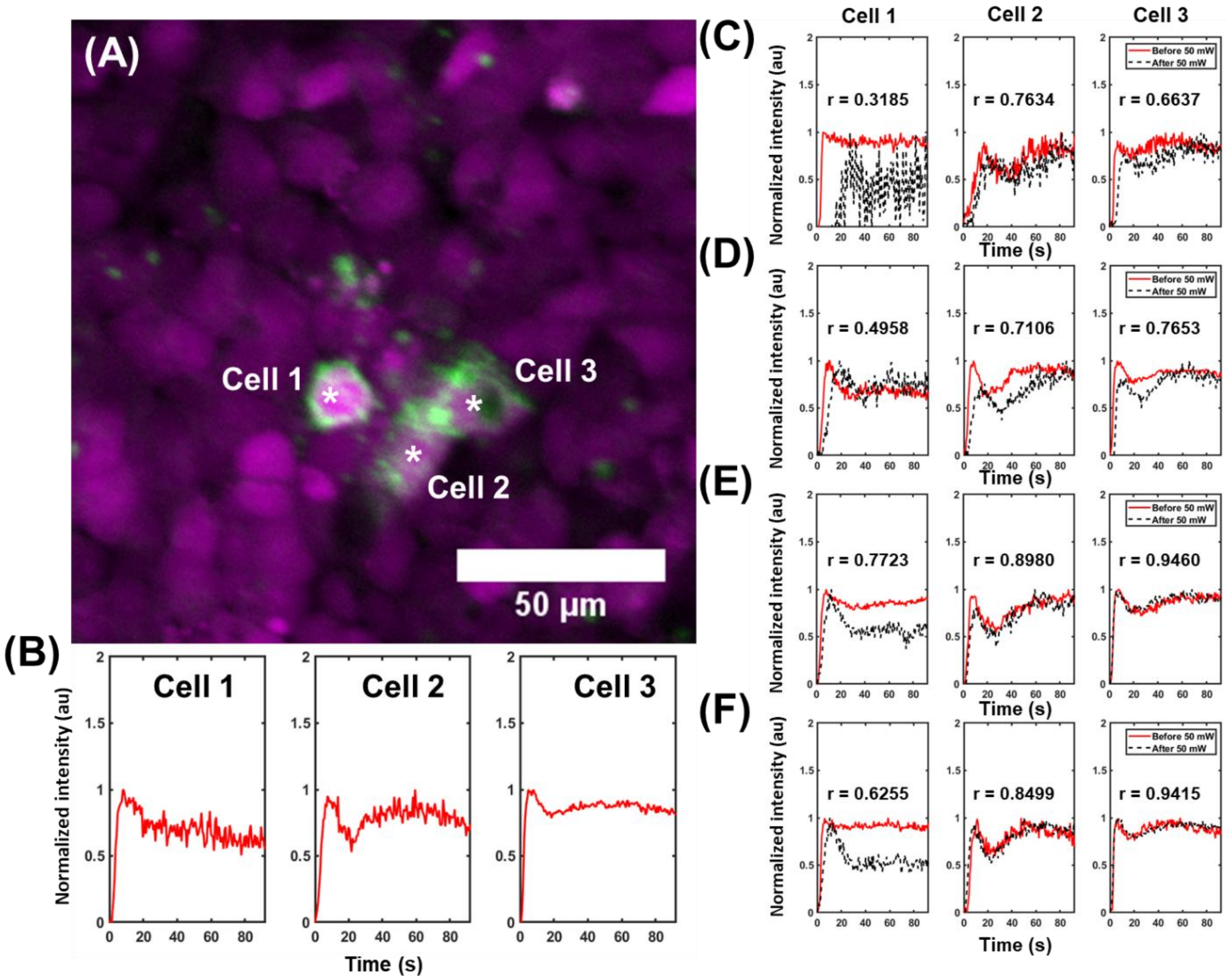


**Supplementary Fig. 13: Power evaluation on melanopsin-expressing HEK293T cells. a,** Representative field-of-view, with Calbryte-590 in magenta and GFP in green. **b,** Calcium transients evoked from Cells 1-3 in **a** at 50 mW, the highest power used in these experiments. These cells were also evaluated at **c,** 5 mW, **d,** 10 mW, **e,** 20 mW, and **f,** 30 mW before (red traces) and after (black traces) illumination with 50 mW. All traces were normalized to 1. Pearson's correlation coefficients are also included in the plots to compare the similarity of profiles before and after illumination with 50 mW. Evaluated in n = 3 cells.

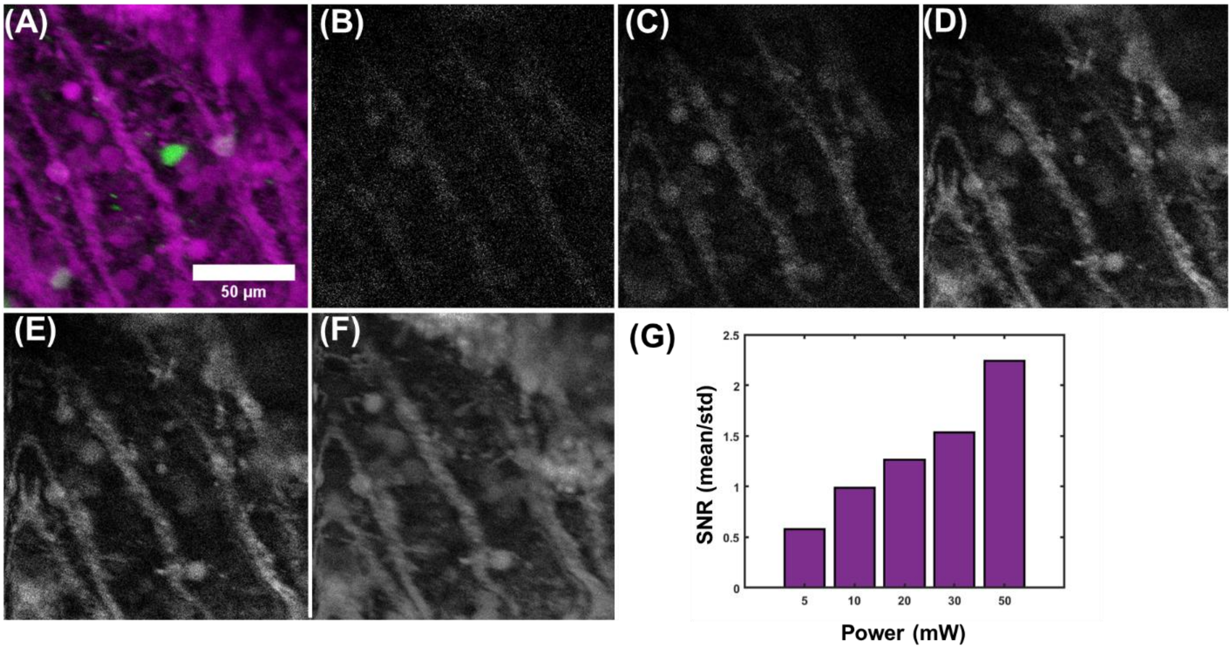


**Supplemental Fig. 14: Evaluation of power on image SNR in retinal tissue. a,** Representative field-of-view for power evaluation. The image for the field-of-view is shown at **b,** 5 mW, **c,** 10 mW, **d,** 20 mW, **e,** 30 mW, **f,** and 50 mW. **g,** Image SNR, calculated as the mean intensity of the image divided by its standard deviation. Results for n = 1 mouse and retina.

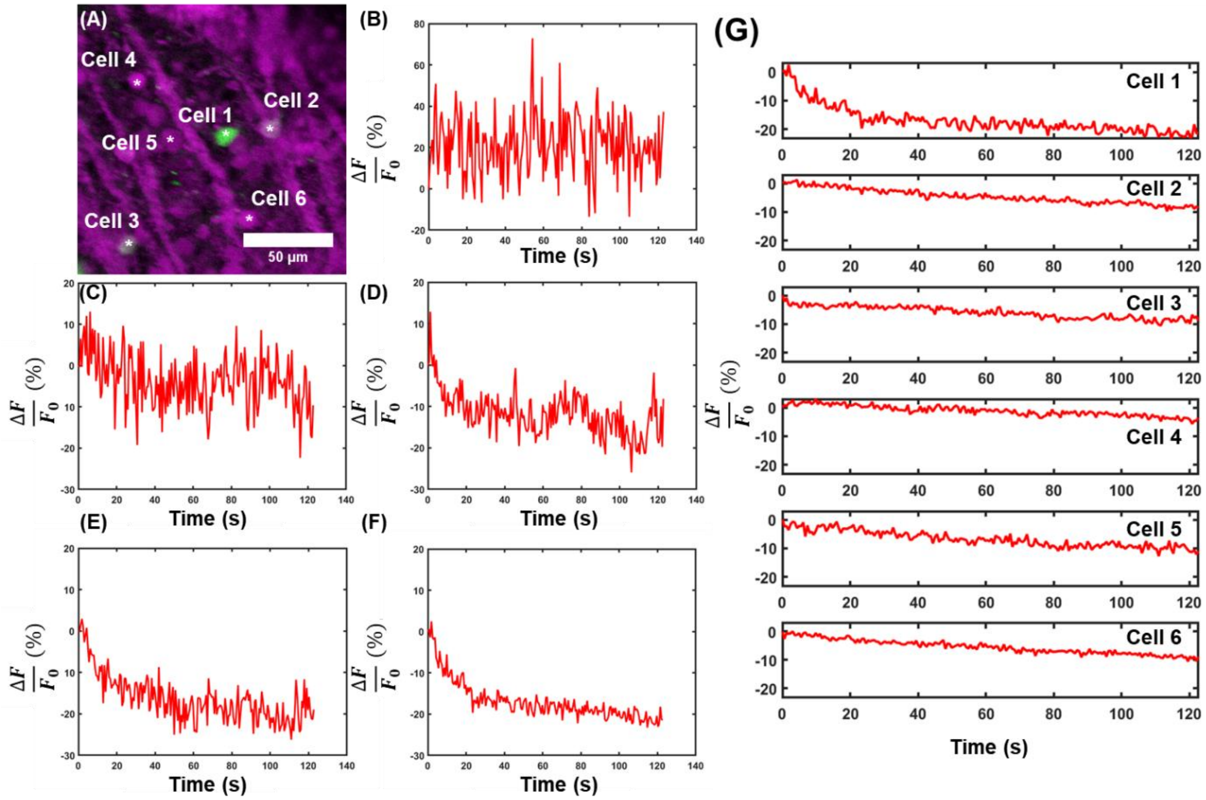


**Supplementary Fig. 15:** Calcium transients for a single field-of-view at increasing powers. **a,** Chosen field-of-view from isolated retina. Calcium transients are illustrated for incident powers at **b,** 5 mW, **c,** 10 mW, **d,** 20 mW, **e,** 30 mW, **f,** and 50 mW. **g,** Transients for all cells in **a**, illuminated with 50 mW. Results from n = 1 cell, N = 1 mouse and retina.
